# Groundwater *Elusimicrobia* are metabolically diverse compared to gut microbiome *Elusimicrobia* and some have a novel nitrogenase paralog

**DOI:** 10.1101/765248

**Authors:** Raphaël Méheust, Cindy J. Castelle, Paula B. Matheus Carnevali, Ibrahim F. Farag, Christine He, Lin-Xing Chen, Yuki Amano, Laura A. Hug, Jillian F. Banfield

**Affiliations:** Department of Earth and Planetary Science, University of California, Berkeley, Berkeley, CA 94720, USA; Innovative Genomics Institute, Berkeley, CA 94720, USA; School of Marine Science and Policy, University of Delaware, Lewes, DE 19968, USA; Nuclear Fuel Cycle Engineering Laboratories, Japan Atomic Energy Agency, Tokai-mura, Ibaraki, Japan; Department of Biology, University of Waterloo, ON, Canada

## Abstract

Currently described members of *Elusimicrobia*, a relatively recently defined phylum, are animal-associated and rely on fermentation. However, free-living *Elusimicrobia* have been detected in sediments, soils and groundwater, raising questions regarding their metabolic capacities and evolutionary relationship to animal-associated species. Here, we analyzed 94 draft-quality, non-redundant genomes, including 30 newly reconstructed genomes, from diverse animal-associated and natural environments. Genomes group into 12 clades, 10 of which previously lacked reference genomes. Groundwater-associated *Elusimicrobia* are predicted to be capable of heterotrophic or autotrophic lifestyles, reliant on oxygen or nitrate/nitrite-dependent respiration, or a variety of organic compounds and *Rhodobacter* nitrogen fixation-dependent (Rnf-dependent) acetogenesis with hydrogen and carbon dioxide as the substrates. Genomes from two clades of groundwater-associated *Elusimicrobia* often encode a new group of nitrogenase paralogs that co-occur with an extensive suite of radical S-Adenosylmethionine (SAM) proteins. We identified similar genomic loci in genomes of bacteria from the Gracilibacteria phylum and the *Myxococcales* order and predict that the gene clusters reduce a tetrapyrrole, possibly to form a novel cofactor. The animal-associated *Elusimicrobia* clades nest phylogenetically within two free-living-associated clades. Thus, we propose an evolutionary trajectory in which some *Elusimicrobia* adapted to animal-associated lifestyles from free-living species via genome reduction.

## Introduction

*Elusimicrobia* is an enigmatic bacterial phylum. The first representatives were termite gut-associated [1], although one 16S ribosomal ribonucleic acid (rRNA) gene sequence was identified in a contaminated aquifer [2]. Initially referred to as Termite Group 1 (TG1) [3], the phylum was renamed as *Elusimicrobia* after isolation of a strain from a beetle larvae gut [4]. *Elusimicrobia* is part of the *Planctomycetes-Verrucomicrobia-Chlamydia* superphylum (PVC). Currently available sequences of *Elusimicrobia* form a monophyletic group distinct from other PVC phyla which include *Planctomycetes, Verrucomicrobia, Chlamydia, Omnitrophica, Desantisbacteria* [5], *Kiritimatiellaeota* [6] and *Lentisphaerae* [7].

The current understanding of *Elusimicrobia* mostly relies on a single taxonomic class for which the name *Endomicrobia* has been proposed [8]. *Endomicrobia* comprise abundant members of the hindgut of termites, cockroaches and other insects, as well as in rumen where they occur as endosymbionts [9], or ectosymbionts [10] of various flagellated protists, or as free-living bacteria [11]. Of the three *Endomicrobia* genomes that have been described, all belong to the genus *Endomicrobium*. One is for a free-living *E*. *proavitum* [12] and two are *E*. *trichonymphae* Rs-D17 endosymbionts, genomovar Ri2008 [13] and genomovar Ti2015 [14]. The fourth *Elusimicrobia* genome available is for *Elusimicrobium minutum* strain Pei191^T^ [15], which was isolated from a beetle larva gut [4]. *E*. *minutum* forms a distinct monophyletic family-level lineage of gut-adapted organisms for which the name *Elusimicrobiaceae* was proposed [4]. Cultivation and genome-based studies revealed that *E*. *proavitum* and *E*. *minutum* strain Pei191^T^ are strictly anaerobic ultramicrobacteria capable of glucose fermentation [4, 16, 17].

Despite prior focus on gut-associated *Elusimicrobia*, phylogenetic analysis of environmental sequences revealed numerous novel and distinct lineages from a wide range of habitat types, including soils, sediments and groundwater [16, 18]. Moreover, several published metagenomics analyses reconstructed genomes from *Elusimicrobia* but none of these studies analyzed them in detail [5, 19, 20]. We augmented the sampling with 30 unpublished genomes from animal-associated and groundwater-associated metagenomes from sequencing efforts that targeted all bacteria and archaea in the metagenomes. Here, we present a comparative genomic analysis of 94 draft-quality and non-redundant genomes from diverse environments. We identified 12 lineages, including 10 that previously lacked genomic representatives. We predict numerous traits that constrain the roles of *Elusimicrobia* in biogeochemical cycles, identify a new nitrogenase paralog, and infer the evolutionary processes involved in adaptation to animal-associated lifestyles.

## Materials and Methods

### Elusimicrobia genome collection

Thirty unpublished genomes were added to the 120 genomes downloaded from the genome NCBI database in June 2018.

Six genomes were obtained from groundwater samples from Genasci Dairy Farm, located in Modesto, California (CA) and from Serpens Ridge, a private property located in Middletown, CA. Over 400 L of groundwater were filtered from monitoring well 5 on Genasci Dairy Farm, located in Modesto, CA, over a period ranging from March 2017 to June 2018. Over 400 L of groundwater were filtered from Serpens Ridge, a private property located in Middletown, CA in November 2017. Deoxyribonucleic acid (DNA) was extracted from all filters using Qiagen DNeasy PowerMax Soil kits and ∼10 Gbp of 150 bp, paired end Illumina reads were obtained for each filter. Scaffolds were binned on the basis of GC content, coverage, presence of ribosomal proteins, presence/copies of single copy genes, tetranucleotide frequency, and patterns of coverage across samples. Bins were obtained using manual binning on ggKbase [21], Maxbin2 [22], CONCOCT [23], Abawaca1, and Abawaca2 (https://github.com/CK7/abawaca), with DAS Tool [24] used to choose the best set of bins from all programs. All bins were manually checked to remove incorrectly assigned scaffolds using ggKbase.

The remaining 23 genomes came from previous sequencing efforts. In brief, three genomes were obtained from sediment samples of Tibet hot springs in 2016. The samples were collected as previously described [25], and for DNA processing and sequencing methods refer to [26]. One genome was obtained from sediment samples collected from Zodletone spring, an anaerobic sulfide and sulfur-rich spring in Western Oklahoma. Detailed site descriptions and metagenomic sequencing were previously reported in [27]. Seven genomes were obtained from groundwater samples collected from the Mizunami and the Horonobe Underground Research Laboratories (URL) in Japan. For sampling refer to [28] and [29], and for DNA processing and sequencing methods refer to [29]. Six genomes were obtained from an aquifer adjacent to the Colorado River near the town of Rifle, Colorado, USA in 2011 [19] one genome from the Crystal Geyser system in Utah, USA [30]. For DNA processing and sequencing methods see [5, 19] Finally, six genomes were assembled from mammal microbiome raw data used in previous studies following the methods described in [31] (**Suppl. Dataset 1**).

### Genome completeness assessment and de-replication

Genome completeness and contamination were estimated based on the presence of single-copy genes (SCGs) as described in [19]. Genome completeness was estimated using 51 SCGs, following [19]. Genomes with completeness > 70% and contamination < 10% (based on duplicated copies of the SCGs) were considered as draft-quality genomes. Genomes were de-replicated using dRep (version v2.0.5 with average nucleotide identity (ANI) > 99%) [32]. The most complete genome per cluster was used in downstream analyses.

### 16S rRNA gene phylogeny

To construct a comprehensive 16S rRNA gene tree, all gene sequences assigned to the *Elusimicrobia* in the SILVA database [33] were exported. Sequences longer than 750 bp were clustered at 97% identity using uclust, and the longest representative gene for each cluster included in the phylogenetic tree. All 16S rRNA genes associated with a binned *Elusimicrobia* genome were added to the SILVA reference set, along with an outgroup of *Planctomycetes, Verrucomicrobia, Chlamydia, Omnitrophica, Desantisbacteria, Kiritimatiellaeota* and *Lentisphaerae* rRNA gene sequences from organisms with published genomes (45 outgroup sequences total). The final dataset comprised 711 sequences, which were aligned together using the SILVA SINA alignment algorithm. Common gaps and positions with less than 3% information were stripped from the alignment, for a final alignment of 1,593 columns. A maximum likelihood phylogeny was inferred using RAxML [34] version 8.2.4 as implemented on the CIPRES high performance computing cluster [35], under the GTR model of evolution and the MRE-based bootstopping criterion. A total of 408 bootstrap replicates were conducted, from which 100 were randomly sampled to determine support values.

### Concatenated 16 ribosomal proteins phylogeny

A maximum-likelihood tree was calculated based on the concatenation of 16 ribosomal proteins (L2, L3, L4, L5, L6, L14, L15, L16, L18, L22, L24, S3, S8, S10, S17, and S19). Homologous protein sequences were aligned using Muscle [36], and alignments refined to remove regions of ambiguity by removing columns with less than 3% information and manually removing abberrent N and C-terminal extensions. The protein alignments were concatenated, with a final alignment of 147 genomes and 2,388 positions. The tree was inferred using RAxML [34] (version 8.2.10) (as implemented on the CIPRES web server [35]), under the LG plus gamma model of evolution, and with the number of bootstraps automatically determined via the MRE-based bootstopping criterion. A total of 108 bootstrap replicates were conducted, from which 100 were randomly sampled to determine support values.

### GTDB taxonomic assignment

Taxonomy assignment was performed for each genome using the GTDB-Tk software (v1.0.1) (default parameters) [37].

### Protein clustering

Protein clustering into families was achieved using a two-step procedure as previously described in [38]. A first protein clustering was done using the fast and sensitive protein sequence searching software MMseqs2 (version 9f493f538d28b1412a2d124614e9d6ee27a55f45) [39]. An all-vs-all search was performed using e-value: 0.001, sensitivity: 7.5 and cover: 0.5. A sequence similarity network was built based on the pairwise similarities and the greedy set cover algorithm from MMseqs2 was performed to define protein subclusters. The resulting subclusters were defined as subfamilies. In order to test for distant homology, we grouped subfamilies into protein families using an HMM-HMM comparison procedure as follows: the proteins of each subfamily with at least two protein members were aligned using the result2msa parameter of mmseqs2, and, from the multiple sequence alignments, HMM profiles were built using the HHpred suite (version 3.0.3) [40]. The subfamilies were then compared to each other using hhblits [41] from the HHpred suite (with parameters -v 0 -p 50 -z 4 -Z 32000 -B 0 -b 0). For subfamilies with probability scores of ≥ 95% and coverage ≥ 0.50, a similarity score (probability X coverage) was used as weights of the input network in the final clustering using the Markov Clustering algorithm [42], with 2.0 as the inflation parameter. These clusters were defined as the protein families.

### Phylogenetic analyses of individual protein sequences

Each individual tree was built as follows. Sequences were aligned using MAFFT (version 7.390) (--auto option) [43]. Alignment was further trimmed using Trimal (version 1.4.22) (--gappyout option) [44]. Tree reconstruction was performed using IQ-TREE (version 1.6.6) (as implemented on the CIPRES web server [35]) [45], using ModelFinder [46] to select the best model of evolution, and with 1000 ultrafast bootstrap [47].

### Protein sequence annotation

Protein sequences were functionally annotated based on the accession of their best Hmmsearch match (version 3.1) (E-value cut-off 0.001) [48] against an HMM database constructed based on ortholog groups defined by the KEGG [49] (downloaded in June 10, 2015). Domains were predicted using the same Hmmsearch procedure against the Pfam database (version 31.0) [50]. The domain architecture of each protein sequence was predicted using the DAMA software (version 1.0) (default parameters) [51]. SIGNALP (version 4.1) (parameters: -f short -t gram-) [52] was used to predict the putative cellular localization of the proteins. Prediction of transmembrane helices in proteins was performed using TMHMM (version 2.0) (default parameters) [53]. The transporters were predicted using BLASTP (version 2.6.0) [54] against the TCDB database (downloaded in February 2019) (keeping the best hit, e-value cut-off 1e-20) [55].

### Metabolic pathways annotation

The Embden-Meyerhof pathway (module M00001), the pentose phosphate pathway (modules M00006 and M00007), cobalamin biosynthesis (M00122) and pyruvate oxidation (module M00307) were considered present based on the completeness of their corresponding KEGG modules (complete if at least 80% of the KEGG module, partial if between 50 and 80%, absent otherwise). The capacity of synthesizing common energy-storage polysaccharides (starch or glycogen) was considered if both the starch/glycogen synthase (KEGG accessions K00703, K20812 or K13679) and 1,4-alpha-glucan branching enzyme were present (K00700 or K16149) [56]. F-type (module M00157, 8 subunits) and V/A-type ATPases (module M00159, 9 subunits) were considered complete if at least 80% of the subunits were present, partial if between 50 and 80%, absent otherwise). The tricarboxylic acid (TCA) cycle was considered complete if 70% of the key enzymes represented by their KEGG accessions were present (partial if between 50 and 70%, absent otherwise). The key enzymes of the TCA cycle include the aconitate hydratase (K01681 or K01682), the fumarate hydratase (K01676 or K01679 or K01677 and K01678), the isocitrate dehydrogenase (K00031 or K00030), the citrate synthase (K01647 or K05942), the malate dehydrogenase (K00024 or K00025 or K00026 or K00116), the succinate dehydrogenase (K00239, K00240 and K00241), the 2-oxoglutarate dehydrogenase (K00174 and K00175) and the succinyl-CoA synthetase (K01899 and K01900 or K01902 and K01903 or K08692 and K01903). The *Rhodobacter* nitrogen fixation (Rnf) complex was considered complete in a genome if at least 4 out of 6 subunits were found in operon along the genome (subunits ABCDEG represented by the KEGG accessions K03617, K03616, K03615, K03614, K03613 and K03612 respectively). The Wood–Ljungdahl (WL) pathway is divided into two branches, the methyl and the carbonyl branches. For the carbonyl branch, reference sequences of the five subunits of the carbon monoxide dehydrogenase/acetyl-coenzyme A (CoA) synthase (CODH/ ACS) were investigated. The AcsE (KPK98995.1 and KPJ61844.1), the CdhA/AcsA (KPK97464.1 and KPJ58813.1), the CdhC/AcsB (KPK97461.1 and KPJ58814.1), the CdhD/AcsD (KPK98994.1 and KPJ61843.1) and the CdhE/AcsC (KPK98991.1 and KPJ61839.1) protein sequences from *Planctomycetes bacterium* DG_23 and *Omnitrophica* WOR_2 bacterium SM23_72 [57] were searched against *Elusimicrobia* to retrieve similar sequences using BLASTP (e-value 1e^−10^) (version 2.6.0+) [54]. The carbonyl branch was considered complete if 4 out of 5 subunits were present (partial if 3 subunits). The methyl branch were considered complete if the alpha subunit of the formate dehydrogenase (NADP+) (K05299), the formate--tetrahydrofolate ligase (K01938), the methylenetetrahydrofolate dehydrogenase (NADP+)/methenyltetrahydrofolate cyclohydrolase (folD) (K01491), methylenetetrahydrofolate reductase (metF, MTHFR) (K00297) or if three key enzymes were detected on the same scaffold (partial if not on the same scaffold). The butyrate fermentation pathway was considered complete in a genome if the two subunits of the acetate CoA/acetoacetate CoA-transferase (K01034 and K01035), the butyrate kinase (K00929) and the butyryl-CoA dehydrogenase (K00248) were present. The acetate metabolism pathway was considered complete if both the acetate/propionate kinase (K00925 or K00932) and the phosphate acetyltransferase (K00625, K13788 or K15024) were present or if the two subunits of the acetate-CoA ligase were present (K01905 and K22224). The ethanol fermentation pathway was considered complete if both the aldehyde dehydrogenase (K00128, K00129, K14085, K00149 or K00138) and the alcohol dehydrogenase (K00114, K13951, K13980, K13952, K13953, K13954, K00001, K00121 or K18857) were present or if the multifunctional aldehyde-alcohol dehydrogenase (encoded by the *adhE* gene, K04072) that catalyzes the sequential reduction of acetyl-CoA to acetaldehyde and then to ethanol under fermentative conditions was present [58]. Lactate fermentation was considered present if the lactate dehydrogenase was present (K00016) and malate fermentation if malate dehydrogenase was present (K00024) [59]. We searched the *Elusimicrobia* genomes for evidence of uroporphyrinogen III synthesis by looking for genes encoding the porphobilinogen synthase (K01698), the hydroxymethylbilane synthase (K01749) and the uroporphyrinogen synthase (K01719). These three enzymes are part of the three-enzymatic-step core pathway from 5-aminolevulinate to uroporphyrinogen III [60]. Based on KEGG accessions, we also annotated the ammonium transporter (K03320), the transhydrogenase (NfnAB) (K00324, K00325), the electron transfer flavoprotein (EtfAB) (K03522, K03521), the ferredoxin nitrite reductase (NirK and NirS) (K00368, K15864), the cytochrome-c nitrite reductase (NfrAH) (K03385, K15876), the phosphotransferase system (PTS) glucose-specific IIC component (K02779), the PTS N-acetylglucosamine-specific IIC component (K02804), the PTS sucrose-specific IIC component (K02810), the PTS maltose/glucose-specific IIB component (K02790), the PTS fructose-specific IIA component (K02768) and the PTS mannose-fructose-sorbose family (K02793, K02794, K02795 and K02796). The succinate dehydrogenase was considered complete in a genome if at least 3 out of 4 subunits were found in operon along the genome (K00239, K00240, K00241, K00242) (KEGG module M00149). The cytochrome *bd* ubiquinol oxidase was considered complete in a genome if the two subunits were found in operon along the genome (K00425 and K00426). The cytochrome *bo*_3_ was considered complete if two subunits out of four were present (K02297, K02298, K02299 and K02300). Hydrogenases sequences were retrieved using Hmmsearch (version 3.1) (E-value cut-off 0.001) [48] and custom HMMs built from previous studies [61, 62]. All hydrogenases were annotated based on the reconstruction of phylogenetic trees and careful inspection of the genes next to the catalytic subunits. A similar strategy was employed to annotate the formate dehydrogenase, the alternative complex III, the nitrite oxidoreductase (NxrA) and the nitrate reductases (NapA, NasA, and NarB) thanks to a phylogenetic tree of the dimethyl sulfoxide (DMSO) reductase superfamily [61, 63] and to annotate the types AAA and cbb cytochrome oxidase and quinol nitric oxide reductase (qNOR) thanks to a phylogenetic tree of the catalytic subunit I of heme-copper oxygen reductases [61].

## Results and Discussion

### Selection and phylogenetic analysis of a non-redundant set of genomes

As of June 2018, 120 genomes assigned to *Elusimicrobia* were available in the NCBI genome database. We augmented these genomes with 30 newly reconstructed genomes from metagenomic samples from Zodletone spring sediments (Oklahoma, USA), groundwater or aquifer sediments (Serpens Ridge, California, USA; Rifle, Colorado, USA; Genasci dairy, Modesto, California, USA; Horonobe, Japan), hot spring sediments (Tibet, China), and the arsenic impacted human gut (see Materials and Methods). Genomes with >70% completeness and <10% contamination were clustered at ≥ 99% ANI, and 113 genomes representative of the clusters were selected based on completeness and contamination metrics. Among them, 19 genomes were missing many ribosomal proteins and thus were excluded, resulting in a final dataset of 94 non-redundant *Elusimicrobia* genomes of good quality (median completeness of 92%, 69 genomes were >90% completeness, **Suppl. Dataset 1**). Of the 94 genomes, 23 are from intestinal habitats and 71 come from other habitats, mostly groundwater-associated (**Suppl. Dataset 1**). Although some groundwater genomes are publicly available [19], none have been analyzed in detail.

The *Elusimicrobia* genomes were further classified into lineages, consistent with the previous classification based on 16S rRNA gene sequences [4, 16]. In order to improve the phylogenetic tree, we also added 16S rRNA gene sequences from genomes that were not originally chosen as representative genomes. The 16S rRNA genes were binned for 41 of the 94 genomes, 31 out of 41 belong to lineages I (*Endomicrobia*), IIa, IIc, III (*Elusimicrobiaceae*), IV, V and VI. Thus, the 16S rRNA gene sequences investigated here span 7 of the 9 previously defined lineages (**Figure S1**) [4, 16]. The remaining 10 sequences do not belong to the well-defined lineages [4, 16] but instead cluster in groups defined by Yarza *et al* [64]. In total, 9 sequences were considered as new phyla by Yarza *et al* whereas the remaining 32 sequences cluster into one single phylum. We compared the classification based on the 16S rRNA sequences with the recent classification proposed by the Genome Taxonomy Database (GTDB) based on concatenated protein phylogeny [65] and found that the 94 genomes are classified into three potentially phylum-level lineages. Of these, 89 genomes genuinely belong to the *Elusimicrobia* phylum (**Suppl. Dataset 1**).

To clarify the taxonomy, we reconstructed a phylogenetic tree based on 16 ribosomal proteins (RPs) and mapped the results of the classifications based on the 16S rRNA and the GTDB (**Figure 1, Figure S2 and Suppl. Dataset 1**). The 94 genomes form a supported monophyletic group (**Figure 1A**, bootstrap: 95) with *Desantisbacteria* and *Omnitrophica* as sibling phyla (**Figure 1A**, bootstrap: 89). The bootstrap support (75) was insufficient to confirm whether *Desantisbacteria* or *Omnitrophica* is the most closely related phylum. All of the gut-associated genomes are clustered into the two known lineages, i.e., *Endomicrobia* (lineage I) and *Elusimicrobiaceae* (lineage III). The *Endomicrobia* lineage formed a distinct clade (n=6, bootstrap: 100) that includes the three earliest described *Endomicrobium* genomes and a new genome from the rumen of sheep [66]. *Elusimicrobiaceae*, which includes genomes related to *Elusimicrobium minutum*, contains genomes from animal habitats, with the exception of one genome from palm oil mill effluent. Of note, two newly reported *Elusimicrobiaceae* genomes in this group are from the gut microbiomes of humans living in Bangladesh [31] and in Tanzania [67]. This is consistent with the recent finding that *Elusimicrobia* genomes are present in the gut microbiomes of non-westernized populations [68]. As previously suggested [4], the lineage III clade (n = 19) is nested within three clades comprising bacteria from groundwater environments (lineages IV, V and VI, n = 38). Based on extrapolated genome sizes, *Elusimicrobiaceae* genomes were significantly smaller than those of sibling lineages IV, V and VI (**Figure S3**). The overall placement and short branch lengths within the *Elusimicrobiaceae* raise the possibility that the *Elusimicrobiaceae* adapted and diversified recently from ancestral groundwater organisms from lineages IV, V and VI. Eleven genomes clustered into lineages IIa (n=2, bootstrap: 41) and IIc (n=9, bootstrap: 100). The lack of bootstrap support does not allow us to confirm the relationship between the two lineages. None of the 94 genomes were classified as lineages IIb and IId, due either to a true absence of IIb and IId genome sequences or because lineages IIb and IId genomes in the dataset lack 16S rRNA gene sequences.

**Figure 1.**
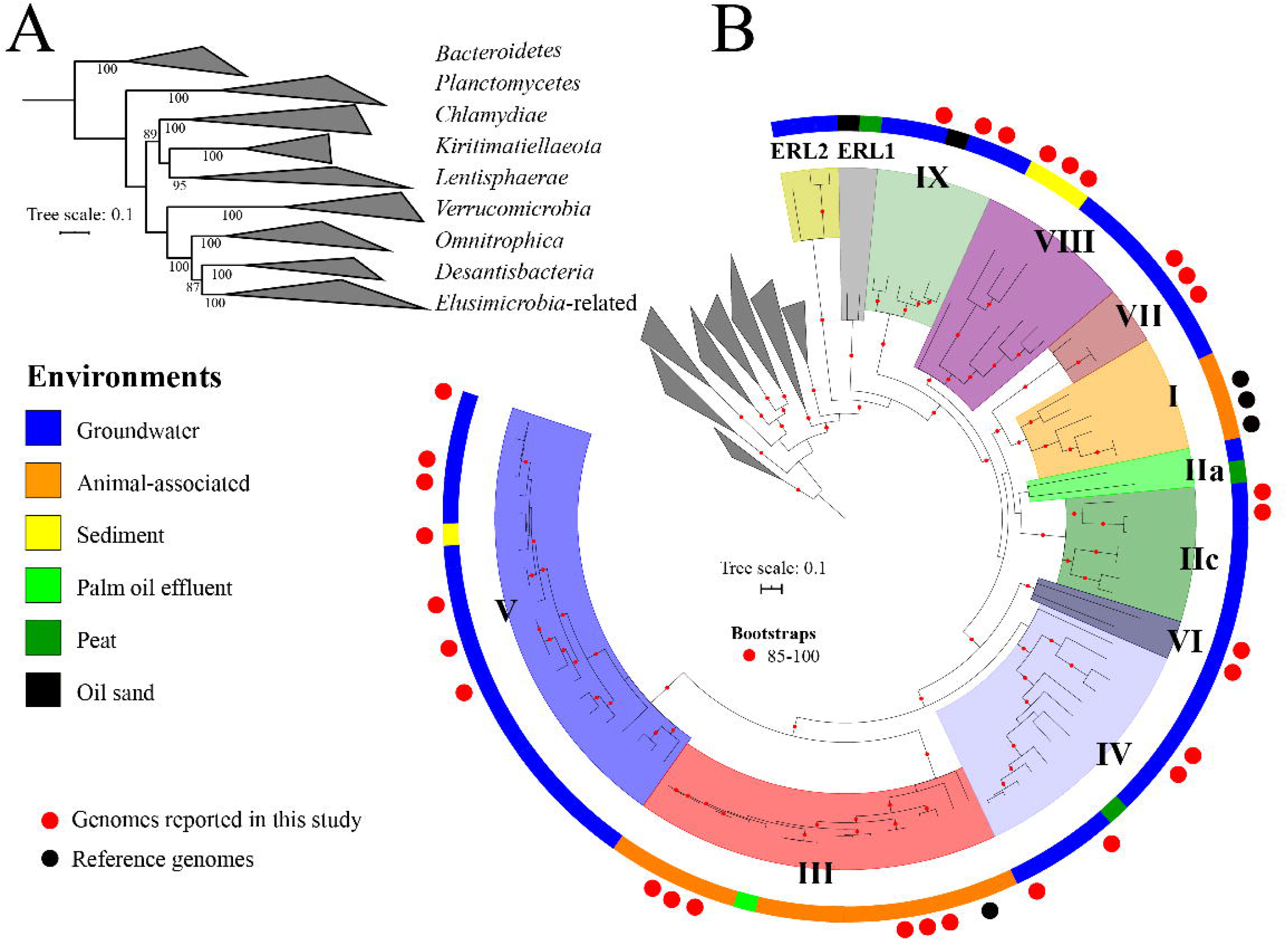
Phylogenetic placement of newly reconstructed genomes. **A**. Relationships between the phyla of the *PVC* superphylum. **B**. Placement of the 94 genomes related to the *Elusimicrobia* phylum. The maximum likelihood tree was constructed based on a concatenated alignment of 16 ribosomal proteins under an LG+gamma model of evolution. The outside color bar indicates the environment of origin. Black circles represent the four genomes described in previous studies while red circles represent the 30 new genomes added in this study. The remaining genomes come from the NCBI genome database. Scale bars indicate the mean number of substitutions per site. The lineages of genomes are indicated by colored ranges and roman numbers. The bootstraps are indicated by red circles when ≥ 85. The complete ribosomal protein tree is available in rectangular format in Figure S2.

The 22 genomes that were not assigned to the previously defined *Elusimicrobia* lineages comprise five clades, of which four are basal to previously reported *Elusimicrobia* lineages. Sixteen of these 22 genomes are from groundwater, and the other 6 genomes were from sediment, oil sand and peat metagenomes. We named three of the five *Elusimicrobia* lineages VII (3 genomes), VIII (8 genomes) and IX (6 genomes), following the current naming procedure. The two other lineages are basal, and are considered as potentially different phyla by both the GTDB and the 16S rRNA gene classification (Figures S1 and S2). These are represented by just five genomes. We assigned these genomes to *Elusimicrobia*-related lineages 1 (ERL1, 2 genomes) and 2 (ERL2, 3 genomes), but the low level of sampling precludes their definitive classification as phyla (**Figure 1 and Figure S2**).

Our analysis captured most of the currently known phylogenetic diversity based on the position in the 16S rRNA gene tree (7 out of the 9 lineages have now a genome). We also discovered 5 new clades based on the ribosomal proteins tree including two phylum-level lineages (**Figure 1**). Many 16S rRNA gene sequences come from soil, especially those from lineages IIa, IIb, IIc, IId and IV (**Figure S1**) [16]. However, the current dataset only includes three genomes from peat environments [20]. Reconstructing genomes from soil is notoriously difficult, as most soils have extremely high microbial diversity [69, 70] and might explain the small number of genomes recovered from this biome.

### A novel nitrogenase paralog possibly involved in the biosynthesis of a novel cofactor

A previous study reported the nitrogen fixing ability of *Endomicrobium proavitum* by an unusual nitrogenase belonging to group IV [17] (**Figure 2**). Dinitrogenase reductase, encoded by *nifH*, donates reducing equivalents to a complex encoded by *nifD* and *nifK*. Canonically, N_2_ fixing NifH proteins belong to groups I, II and III [71]. Analysis of the amino acids coordinating the P-cluster and FeMoCo ligands in NifD indicates that the homologs in the subcluster IVa are functional reductases. Together with the documentation of diazotrophy in *E*. *proavitum*, there is no doubt that some NifH paralogs of group IV comprise functional nitrogenases [17]. Other group IV NifH proteins, non affiliated to the subcluster IVa, are implicated in the biosynthesis of cofactor F430, the prosthetic group of methyl coenzyme M reductase, which catalyzes methane release in the final step of methanogenesis [72, 73]. Another NifH paralog, phylogenetically defined as group V, is involved in chlorophyll biosynthesis [74]. In this case, protochlorophyllide is converted to chlorophyllide via the BchLNB complex in which BchL is the NifH paralog and BchN and BchB are the NifD and NifK paralogs, respectively.

**Figure 2.**
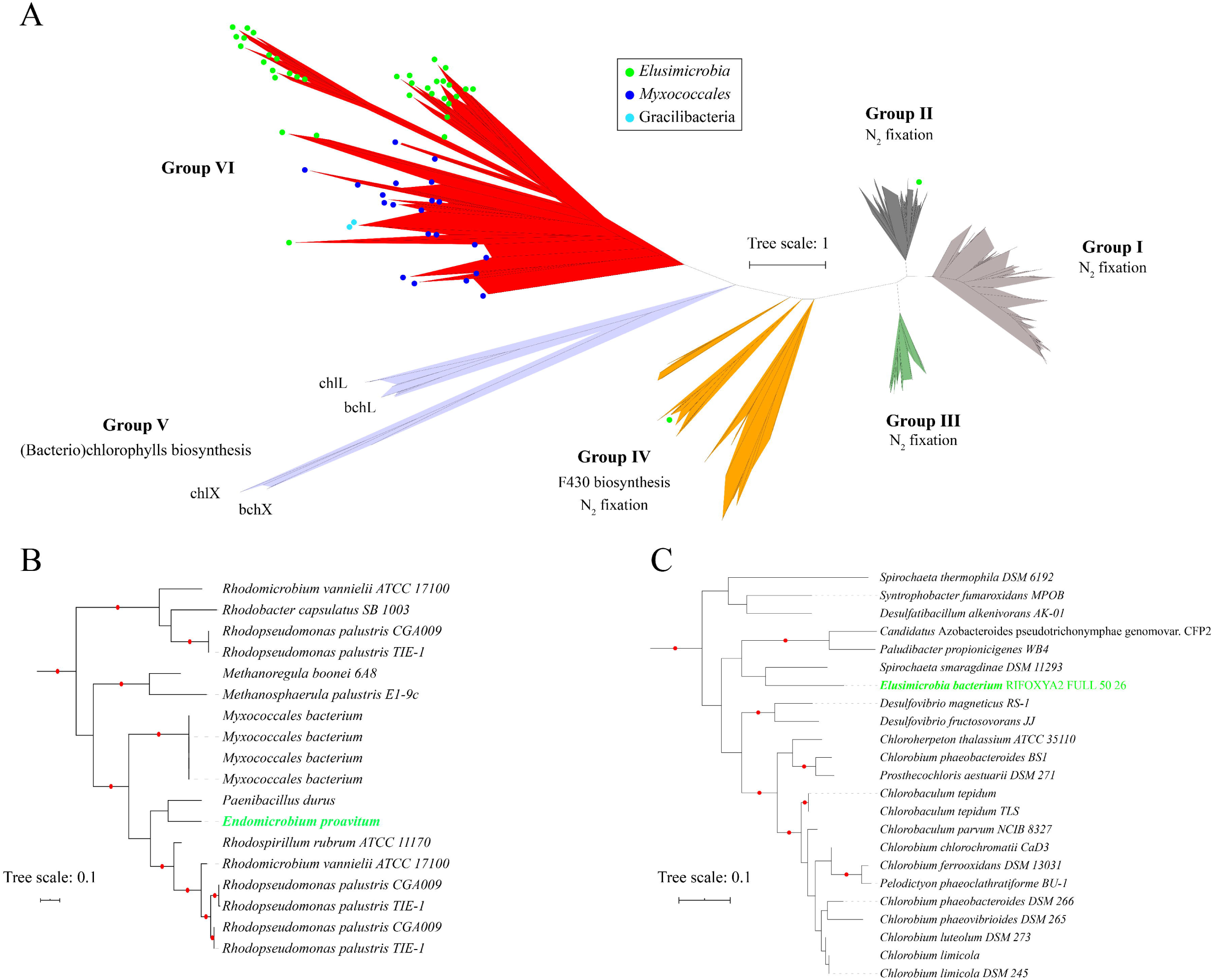
Maximum likelihood phylogeny of the nitrogenase subunit NifH. **A**. Full phylogeny of the NifH subunit. **B**. Detailed phylogeny of the group IV. **C**. Detailed phylogeny of the group II. The tree was inferred using an LG+I+G4 model of evolution. The tree was made using reference sequences from [71]. The green circles highlight the sequences present in *Elusimicrobia* genomes. Scale bar indicates the mean number of substitutions per site. The sequences and tree are available with full bootstrap values in Fasta and Newick format in the Supplementary Data. Branches with bootstrap values ≥ 95 are indicated by red circles.

We searched the *Elusimicrobia* genomes for *nifH, nifD* and *nifK* and identified homologues in 24 *Elusimicrobia* genomes from lineages I, IV and V. We also searched for accessory genes required for the assembly of the M- and P-metallo clusters required by the nitrogenase complex. Two genomes from lineage I, RIFOXYA2_FULL_50_26 and the previously reported *E*. *proavitum*, contain both the nitrogenase subunits and the accessory proteins. The NifH protein from RIFOXYA2_FULL_50_26 places phylogenetically into NifH group II, and thus is plausibly associated with N_2_ fixation [71].

The other NifH proteins all belong to lineages IV and V of *Elusimicrobia* and place phylogenetically outside of all previously described groups of NifH (**Figure 2**). Instead, the NifH sequences from 22 *Elusimicrobia* genomes form a new group that is sibling to group V (**Figure 2**). Consistent with the new placement, these 22 *nifH*-encoding genomes lack the accessory genes required for the assembly of the M- and P-metallo clusters of nitrogenase. We designate these genes as a new group, group VI, and infer that these paralogous proteins may have a distinct and perhaps previously undescribed biological role. The group VI NifH proteins contain the GXGXXG consensus motif for the binding of MgATP and two cysteine (Cys) residues (Cys^97^ and Cys^132^) that bridge the two subunits through a [Fe_4_S_4_] cluster (**Figure S4**). However, the associated *nifD-* and *nifK-like* genes are highly divergent from true nitrogenase genes (25.51 and 22.74 amino acid percent identity on average respectively) (**Figure S5**). Importantly, they lack some conserved cysteine motifs that are involved in the attachment of the P-clusters [71] (**Figure S5**).

Interestingly, nitrogenase paralogs complexes from group IV and V each modify a tetrapyrrole molecule by reducing a carbon-carbon double bound [72, 74, 75]. Biosynthesis of cofactor F430 involves the sirohydrochlorin precursor and biosynthesis of chlorophyll involves a protoporphyrin precursor, both of which derive from uroporphyrinogen III (also a building block for cobalamin). In a subset of genomes with the novel group IV *nifH* genes (five genomes of lineage IV), we identified the capacity to produce uroporphyrinogen III (for example, see *Elusimicrobia* bacterium GWA2_69_24). The absence of precursor biosynthesis pathways in other genomes of *Elusimicrobia* predicted to have the capacity to make nitrogenase clusters does not rule out an analogous function, as many bacteria scavenge such molecules (e.g., cobalamin; [76] or haem [77]).

We examined the genomic neighborhoods to get insights regarding the function of the novel group VI paralogs. The lineage IV genomes encode several copies of *nifK, nifD* and *nifH* whereas most genomes of lineage V genomes have only one copy of each subunit (**Figure S6**). Strikingly, many adjacent genes encode radical S-adenosylmethionine (SAM) proteins. Radical SAM proteins have many functions, including catalysis of methylation, isomerization, sulfur insertion, ring formation, anaerobic oxidation, and protein radical formation, and also in the biosynthesis of cofactors, DNA and antibiotics [78, 79]. The copy number of radical SAM genes varies greatly across the genomes, from no radical SAM genes to 13 copies in close proximity to the nitrogenase paralogs in the genome of GWC2_Elusimicrobia_56_31 (**Figure S6**). Several radical SAM genes are fused with B12-binding domains (SR-2 Biohub_180515 Elusimicrobia_69_71, GWC2_Elusimicrobia_56_31 and *Elusimicrobia bacterium* GWF2_62_30) and/or HemN_C domain (*Elusimicrobia bacterium* GWA2 _69_24) (**Figure S6**). The B12-binding domain is involved in binding cobalamin while the HemN_C domain has been suggested to bind the substrate of coproporphyrinogen III oxidase [80]. Both substrates have tetrapyrrole structures, which is consistent with the phylogenetic position of the NifH near the groups IV and V NifH clades (**Figure 2**). Finally, we constructed a Hidden Markov Model using the new group VI *Elusimicrobia* NifH sequences and searched for homologous sequences in genomes from other phyla. Interestingly, we found homologs of the group VI NifH in two genomes of Gracilibacteria, a phylum within the bacterial Candidate Phyla Radiation [81], and in fourteen *myxococcales* genomes from aquatic environments (**Figure 2 and Suppl. Dataset 2**). Together, these observations suggest that the enzymes likely do not perform nitrogen fixation, but have an alternative function that may be related to the biosynthesis of a tetrapyrrole cofactor similar to chlorophyll or F430 cofactors.

### Overall metabolic potential in *Elusimicrobia* and related genomes

The availability of good quality genomes allowed us to compare their gene contents. We clustered the protein sequences for all genomes to generate groups of homologous proteins (protein families) in a two-step procedure (see Materials and Methods) (**Figure S7**) [38]. The objective was to compare the proteomes across the genomes. Our approach is agnostic and unbiased by preconceptions about the importance of genes, and allows us to cluster protein sequences with no homology within annotation databases. This resulted in 6,608 protein families that were present in at least three distinct genomes. We constructed an array of the genomes versus the protein families, and sorted the families based on profiles of genome presence/absence. Numerous protein families group together due to co-existence in multiple genomes (**Figure S7**). Several blocks of families are fairly lineage specific. This is particularly apparent for the gut-associated *Elusimicrobiaceae* lineage (lineage III) which lacks numerous families abundant in other lineages but also contains 161 families abundant in *Elusimicrobiaceae* but rare or absent in other lineages (red boxes in **Figure S7**). Unlike lineage III, lineage V encodes extended sets of protein families (blue box in **Figure S7**) consistent with their larger genome size (**Figure S3**). Other lineages also have enriched groups of families, although it is less apparent than for lineages III and V (**Figure S7**). Overall, the patterns of presence/absence of protein families are consistent with the lineages defined by the ribosomal protein phylogeny (e.g., genomes from the same lineage tend to have a similar protein families set) and may reflect different metabolic strategies.

Previously represented by gut-associated lineages, the expansion of *Elusimicrobia* identifies a broad variety of energetic strategies within the phylum. Non-gut-associated *Elusimicrobia* display substantial metabolic versatility, with the genomic potential for both autotrophic and heterotrophic-based lifestyles, though not usually both within the same genome or lineage (**Figure 3**). Most of the non-gut-associated genomes are predicted to have the capacity for sugar fermentation to acetate, malate, butyrate and/or ethanol, generating ATP via substrate-level phosphorylation (**Suppl. Dataset 3**). We also identified a large variety of membrane-bound and cytoplasmic [NiFe] and [FeFe] hydrogenases (**Suppl. Dataset 3**), some of which may be electron bifurcating. Other electron bifurcating complexes, such as the transhydrogenase (NfnAB) or the Bcd/EtfAB complex are also present in these genomes (**Suppl. Dataset 3**), suggesting the ability to minimize free energy waste and to optimize energy conservation [82]. This large repertoire of metabolic capacities differs from their animal habitat-residing counterparts (lineages I and III), which rely solely on fermentation for energy [12, 13, 15, 17] (**Figure 3**).

**Figure 3.**
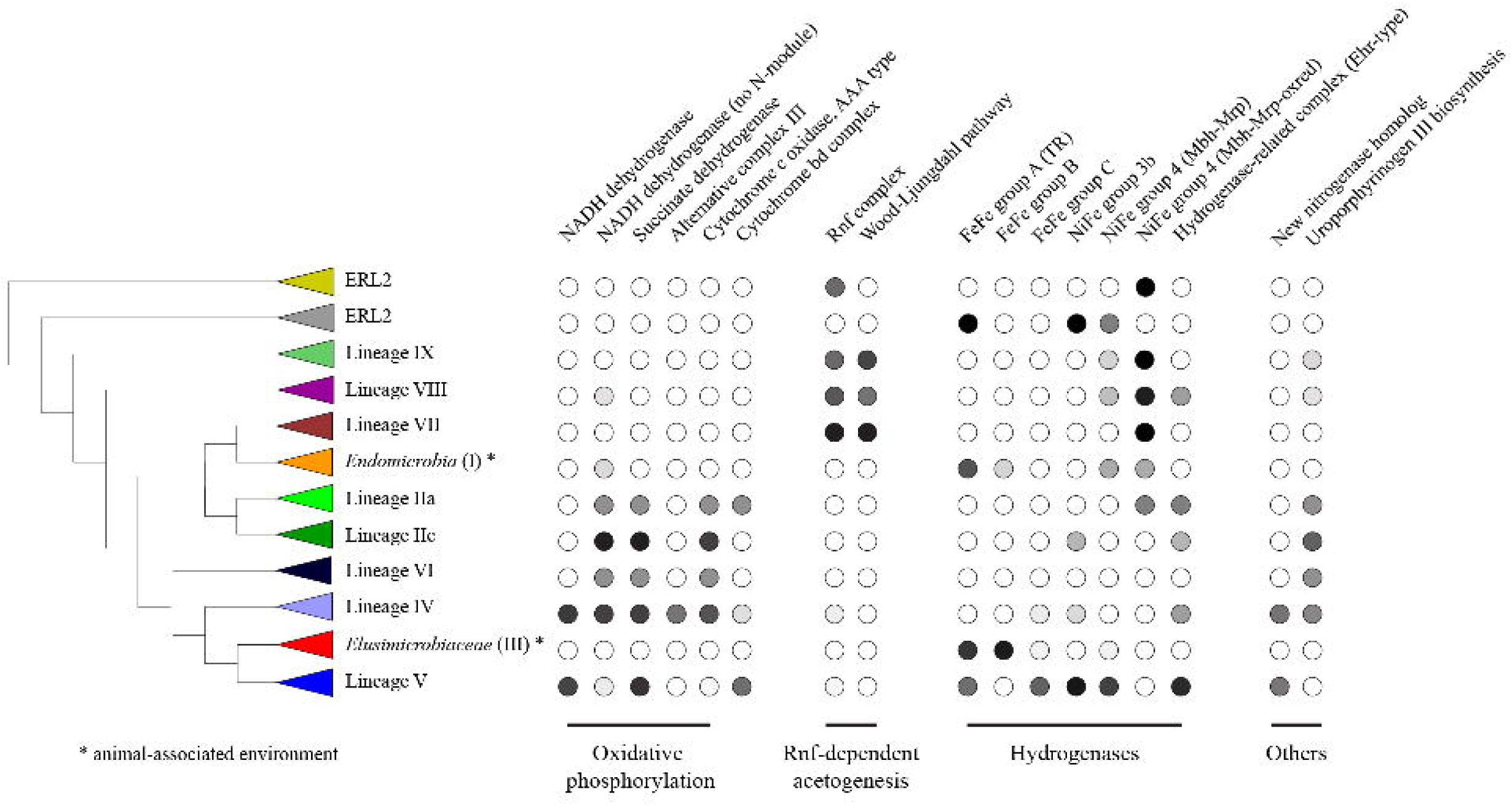
Schematic representation of the relationship of the *Elusimicrobia*-related lineages and the distribution of selected metabolic features across the different members of the phylum. Grey scale of the filled circles represents the prevalence of a function within a given lineage, with black = 100% of genomes and white = 0% of genomes. See **Suppl. Dataset 3** for a fully detailed version of each genome.

### *Elusimicrobia*-related lineages 1 and 2

Analyses of the five genomes assigned to ERL1 and ERL2 (**Figure 1**) suggest that they are likely obligate fermenters. All of the genomes lack a complete tricarboxylic acid cycle, NADH dehydrogenase (complex I), and most other complexes from the oxidative electron transport phosphorylation chain (e.g., complex II, complex III, complex IV, and quinones). Interestingly, all but one genome has partial ATP synthase, so the functionality of this complex V remains uncertain. All genomes included in this study have complete or near-complete glycolysis and/or pentose phosphate pathway(s) and are predicted to have the capacity to produce acetate, lactate, and/or hydrogen as byproducts of fermentation (**Suppl. Dataset 3**). Acetate kinase and phosphotransacetylase have been identified and are likely involved in acetate and ATP production. Thus, we cannot establish that they have a functional complete ATPase, they should be able to produce ATP via substrate level phosphorylation. These organisms may also be capable of synthesizing common energy-storage polysaccharides, as we identified several genes encoding enzymes for starch or glycogen metabolism (**Suppl. Dataset 3**). Overall, we predict widespread fermentation-based metabolism in the nearest neighbour lineages of *Elusimicrobia*.

### Diverse respiratory strategies in groundwater-associated *Elusimicrobia*

Oxidative phosphorylation and the tricarboxylic acid cycle are commonly encoded in genomes from lineages IIa, IIc, IV, V and VI (**Figure 4A**). Lineage V genomes encode a canonical NADH dehydrogenase (complex I with 14 subunits) and succinate dehydrogenase (complex II) that link electron transport to oxygen as a terminal electron acceptor via the high-affinity oxygen cytochrome *bd* oxidase (complex IV). Unlike genomes encoding electron transport chains, lineage V genomes consistently lack a complex III (*bc*_1_-complex, cytochrome c reductase). Organisms that carry one or more bd-type oxidases usually also possess at least one heme-copper oxygen reductase [83]. For instance, *Escherichia coli* also lacks complex III and/or alternative complex III but carries a cytochrome *bo*_3_, a proton-pumping oxidoreductase [83, 84]. However, we did not detect genes encoding cytochrome *bo*_3_ in genomes of lineage V (**Suppl. Dataset 3**). The absence of any heme–copper oxygen reductase was also observed in organisms such as *Lactobacillus plantarum* [85] and *Zymomonas mobilis* [86]. As all of these organisms are capable of oxygen-based respiration, we conclude that there may be a variety of ways to circumvent the lack of complex III (or alternative complex III). Thus, we cannot rule out the possibility that these *Elusimicrobia* have a functional aerobic electron transport chain.

**Figure 4.**
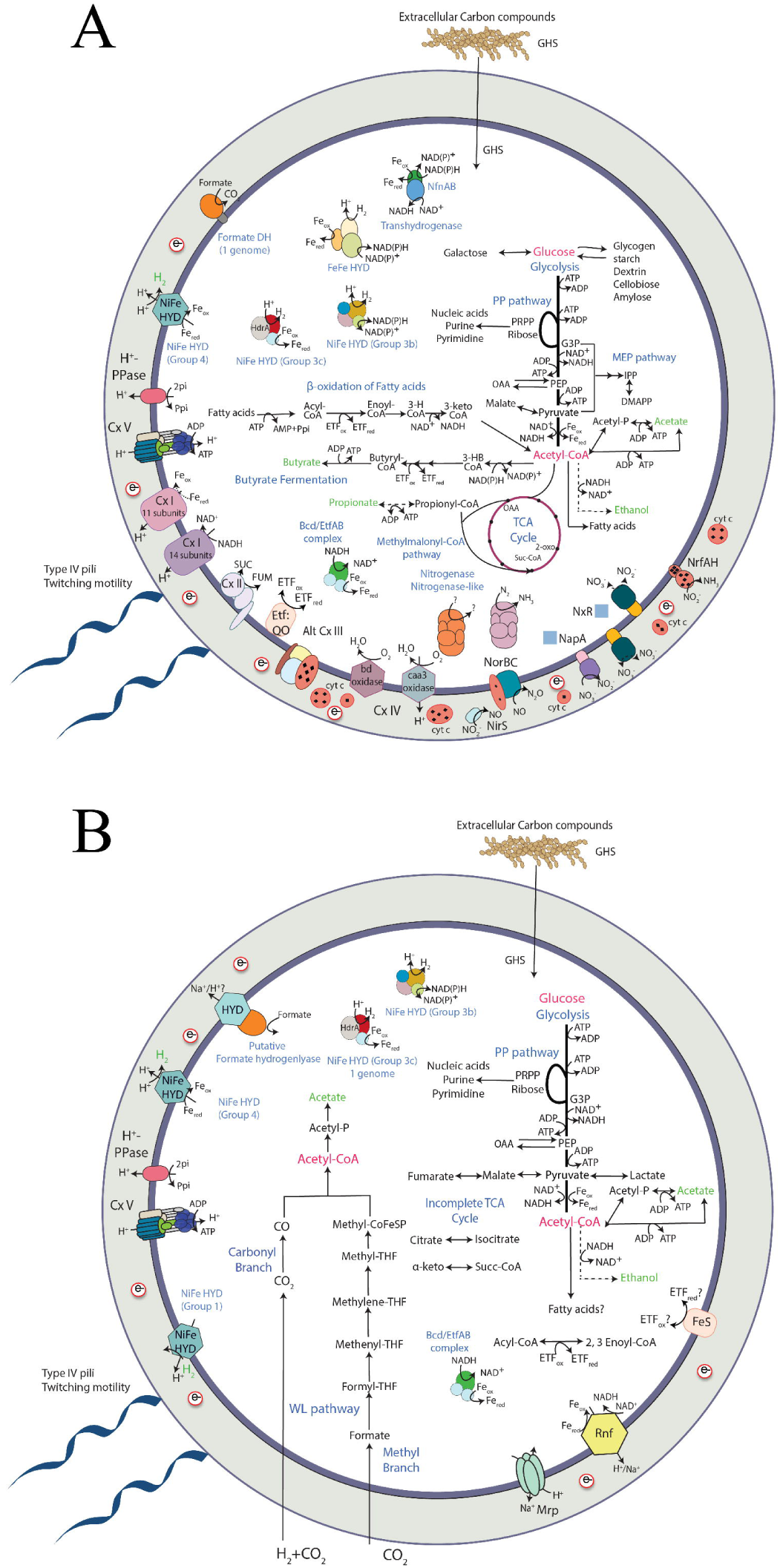
Overview of the metabolic capabilities of non-gut-associated *Elusimicrobia*. A. Lineage IV and V, (B). Lineage IX. Abbreviations not defined in the text: GHS, glycoside hydrolases; TCA, tricarboxylic acid cycle; 2-oxo, 2-oxoglutarate; Suc-CoA, succinyl-CoA; DH, dehydrogenase; HYD, hydrogenase; PPi, pyrophosphate; Pi, inorganic phosphate; PRPP, Phosphoribosyl pyrophosphate; G3P, glyceraldehyde-3-phosphate; PEP, phosphoenolpyruvate; OAA, oxaloacetate; MEP pathway, 2-*C*-methyl-D-erythritol 4-phosphate/1-deoxy-D-xylulose 5-phosphate pathway; IPP, isopentenyl pyrophosphate; DMAPP, dimethylallylpyrophosphate; FeS, FeS oxidoreductase; ETF, electron transfer flavoprotein; ETF:QO, Electron-transferring flavoprotein dehydrogenase/ubiquinone oxidoreductase; BCD/ETFAB, butyryl-CoA dehydrogenase/electron-transfer flavoprotein complex; 3HB-CoA, 3-hydroxybutyryl-CoA dehydrogenase; 3-H-CoA, 3-hydroxyacyl-CoA; 3-keto-CoA, 3-ketoacyl-CoA; H+ – PPase, proton-translocating pyrophosphatase; Cx V, ATP synthase; Cx I, NADH dehydrogenase; Cx II, succinate dehydrogenase/fumarate reductase; Alt Cx III, alternative complex III; Cx IV, cytochrome c oxidase or cytochrome bd oxidase; NorBC, nitric-oxide reductase; NirS, cytochrome cd1-type nitrite reductase; cyt c, c-type cytochrome; NapA, periplasmic nitrate reductase; NxR, nitrite/nitrate oxidoreductase; NrfAH, nitrite reductase; Mrp, multi-subunit Na^+^/H^+^ antiporter; WL pathway, Wood– Ljungdahl pathway; THF, tetrahydrofolate; Rnf, Ferredoxin:NAD+-Oxidoreductase.

Several genomes from lineages IIa, IIc, IV, VI and VI have the genomic potential for respiring a variety of organic compounds (including ribose, galactose, glucose, acetate and possibly propionate and butyrate) as energy and carbon sources. Further, the genomes indicate the capacity for utilization of fatty acids via the ß-oxidation pathway. Examination of the genomes revealed the presence of glycosyl hydrolases that would support growth by utilization of externally derived sugar polymers. Indeed, the organisms from lineages IIa, IIc, IV, VI and VI may also be capable of synthesizing and/or utilizing common energy-storage polysaccharides, as we identified several genes encoding enzymes for starch, glycogen, dextrin metabolism (**Figure 4A**).

Lineage V *Elusimicrobia* typically have a suite of hydrogenases, where hydrogenases are sparsely distributed in other lineages. Some groundwater-associated lineage V genomes have trimeric group A [FeFe] hydrogenases directly downstream from a monomeric group C hydrogenase, related to those seen in *Ignavibacterium album* and *Caldithrix abyssi* [62] (**Figure S8**). In general, [FeFe] hydrogenases can either use or produce H_2_ whereas group C hydrogenases are co-transcribed with regulatory elements and are predicted to sense H_2 [87]_. In most group C hydrogenases this sensing occurs via a Per-ARNT-Sim (PAS) domain [62, 88]. However, the *Elusimicrobia* hydrogenase seems to be fused to a histidine kinase HATPase domain, probably fulfilling this sensory function. Additionally, the genomic neighborhood of these hydrogenases includes several regulatory genes encoding proteins such as Rex, which is known to sense NADH to NAD^+^ ratios for transcriptional regulation [89]. Groundwater-associated genomes of lineage V also encode genes for different types of [NiFe] hydrogenases. Seven genomes encode cytoplasmic group 3b (NADP-reducing) [NiFe] hydrogenases that are likely bidirectional (also known as sulfhydrogenase). Four other genomes also have cytoplasmic group 3c (methyl viologen-reducing) [NiFe] hydrogenases (seen in one other genome in this clade), probably involved in H_2_ utilization (**Figure S9**) [90, 91]. Additionally, most genomes in lineage V encode membrane-bound group 4 [NiFe] hydrogenases of the Mbh-Mrp type, likely involved in H_2_ production (**Figure S10**). Membrane-bound hydrogenases are known to oxidize reduced ferredoxin, and the presence of antiporter-like subunits suggests that they may be involved in ion translocation across the membrane and the generation of a membrane potential [92]. These genomes also encode hydrogenase-related complexes (Ehr), the role of which is still unknown, although it has been suggested that they may interact with oxidoreductases and quinones and may be involved in ion translocation across the membrane [61, 93]. Taken together, this indicates lineage V *Elusimicrobia* sense H_2_ and NADH levels and regulate their metabolisms in response. The repertoire of functions indicated by their suite of hydrogenases suggests they are important players in the hydrogen economy of their ecosystem.

Groundwater and peat-associated lineages IIa, IIc, IV, and VI have somewhat distinct respiratory capacities compared to the groundwater-associated lineage V *Elusimicrobia*. All have a complex I lacking the diaphorase N-module (*nuoEFG* genes), a complex which is hypothesized to use reduced ferredoxin instead of NADH [94]. Lineage IV genomes also have a canonical complex I with an N-module. All genomes from these lineages have complex II (succinate dehydrogenase) and complex IV (cytochrome c oxidase, type A; **Figure S11**). Lineage IV is further different in that it encodes an alternative complex III and a few lineage IV genomes contain a canonical complex III (cytochrome c reductase). Unlike lineage V, which has a consistent set of hydrogenases, some of these genomes have hydrogenases and others do not (**Suppl. Dataset 3**).

The genomes of some members of the groundwater and peat-associated lineages IIa, IIc, IV, V and VI encode nitrogen cycling capacities that are rare in other lineages. Specifically, seven genomes encode a nitrite oxidoreductase (NxrA, **Figure S12**) and/or nitrate reductase (**Figure S12**), indicating that these organisms can respire using nitrate. We also identified other enzymes involved in the nitrogen cycle, such as quinol-dependent nitric-oxide reductase (qNOR, **Figure S11**), cytochrome-c nitrite reductase (NfrAH), and copper and/or cytochrome cd1 nitrite reductase(s) (NirK and NirS) (**Suppl. Dataset 3**). These findings expand the known lifestyles in the phylum *Elusimicrobia* from exclusively fermentative to include several respiratory strategies, using oxygen or nitrogen compounds as terminal electron acceptors for energy conservation.

### RNF-dependent acetogenesis in groundwater-associated *Elusimicrobia*

Other groundwater-associated *Elusimicrobia* in lineages VII, VIII and IX lack the capacity to reduce oxygen or nitrate. Instead, lineages VII, VIII and IX encode the Wood-Ljungdahl pathway (WLP) for the reduction of carbon dioxide to acetyl-coenzyme A with concomitant energy conservation (**Figure 4B and Suppl. Dataset 3**). The WLP is often coupled with cytochromes and quinones to generate a membrane potential for ATP synthesis via an ATP synthase [95]. However, cytochromes and quinones are missing in these lineages. Instead, these genomes encode the Rnf complex, a sodium-motive ferredoxin:NAD oxidoreductase. The Rnf complex generates a sodium ion potential across the cell membrane in *Acetobacterium woodii* [96]. These observations suggest that lineages VII, VIII and IX are capable of autotrophic growth with molecular hydrogen and carbon dioxide as substrates. Members of lineages VII, VIII, IX, ERL2, and a few genomes in lineage I have membrane-bound group 4 [NiFe] hydrogenases of the Mbh-Mrp-oxidoreductase type (**Figure S10 and Suppl. Dataset 3**). The function of this oxidoreductase is unknown, though it has been seen before in other organisms such as the *Thermococcales* [92] and *Saganbacteria* [61]. Both Rnf complexes and group 4 NiFe membrane-bound hydrogenases could extrude ions across the membrane and contribute to the creation of a membrane potential. These *Elusimicrobia* lineages are predicted acetogens, another lifestyle not previously associated with the phylum.

### Gut-associated *Elusimicrobia*

The genomes and general metabolic characteristics of gut-related *Elusimicrobia* (lineages I and III) have been well described previously [12–15]. However, here we extend prior work by detailing the hydrogenases of these bacteria (**Figure 3 and Suppl. Dataset 3**). In brief, prior work indicates that gut-associated *Elusimicrobia* rely on fermentation [4, 12, 13, 16, 17]. Most gut-related *Elusimicrobia* have ‘ancestral’ group B and group A [FeFe] hydrogenases. The ‘ancestral’ group B is typically found in anaerobic bacteria (*e*.*g*., *Clostridia, Actinobacteria*, and *Bacteroidetes*) that inhabit gastrointestinal tracts and anoxic soils and sediments [62, 88] (**Figure S8**). Group B [FeFe] hydrogenases remain largely uncharacterized. It has been suggested that they could couple ferredoxin oxidation to H_2_ evolution, because fermentative organisms rely on ferredoxin as a key redox equivalent [62]. In general, group B enzymes are monomeric. Many of the gut-associated *Elusimicrobia* genomes also harbor a trimeric group A [FeFe] hydrogenase that may be involved in fermentative metabolism. Group A [FeFe] hydrogenases contain NADH-binding domains in the beta subunit and are known to be involved in electron bifurcation or the reverse reaction, electron confurcation (**Figure S8**). In the case of electron bifurcation, [FeFe] hydrogenases oxidize H_2_ by coupling the exergonic reduction of NAD^+^ with the endergonic reduction of oxidized ferredoxin. In electron confurcation, these [FeFe] hydrogenases produce H_2_ by using the exergonic oxidation of ferredoxin with protons (H^+^) to drive the endergonic oxidation of NADH with H^+^ [97–99]. We conclude that many gut-associated *Elusimicrobia* can potentially both produce and use H_2_, but using different hydrogenase types than those of other *Elusimicrobia* lineages.

Distinct from these, two gut-associated *Elusimicrobia* have additional mechanisms to produce H_2_. One genome (genome 277) harbors a monomeric (as opposed to trimeric, as described above) group A [FeFe] hydrogenase in addition to a group B hydrogenase (**Figure S8**). Monomeric group A hydrogenases are thought to be involved in H_2_ production from ferredoxin and flavodoxin [88, 100]. Another gut-associated genome (*Endomicrobium proavitum*) has a tetrameric group A [FeFe] hydrogenase, which is also known to be involved in electron bifurcation and likely H_2_ production [88].

The *Elusimicrobium minutum* Pei191 genome is unusual because it harbors four different kinds of [FeFe] hydrogenases (**Figure S8**), three of which have been described before, a monomeric group B [FeFe] hydrogenase (HydA2), downstream from a dimeric ‘sensory’ group C (HydS) [FeFe] hydrogenase [89] and a trimeric group A [FeFe] hydrogenase [15]. The fourth [FeFe] hydrogenase is another monomeric group B hydrogenase like others found in gut-associated *Elusimicrobia*, with a slightly different domain composition of the catalytic subunit than the other group B hydrogenase (HydA2) in this genome. A membrane-bound group 4 [NiFe] hydrogenase of the Mbh-Mrp type also occurs in this genome [15] (**Figure S10**). This is striking, as the rest of the gut-associated *Elusimicrobia* completely lack [NiFe] hydrogenases (**Suppl. Dataset 3**).

## Conclusions

Previously, our understanding of the *Elusimicrobia* phylum mostly relied on gut-associated habitats although 16S rRNA gene sequences indicated their wider environmental distribution and some genomes from non-gut habitats were available in genome databases. We expanded the genomic representation of this phylum with thirty new genome sequences, mainly from groundwater. However, six of our draft genomes came from humans and pigs, thus expand on the finding of *Elusimicrobia* of lineage III in the gut microbiome of non-Westernized humans [68]. Our results reshape our view of the *Elusimicrobia* phylum, as most of these genomes likely derive from free-living organisms that are predicted to have greater metabolic flexibility than the strictly fermentative prior representatives of the phylum. Our phylogenetic results reveal five new lineages including two potential phylum-level lineages and suggest that animal gut-associated *Elusimicrobia* of lineages I and III evolved from two distinct groups of free-living ancestors, lineages IV, V and group VII, respectively. Based on extant members of groundwater groups, the ancestors of gut-associated *Elusimicrobia* likely underwent substantial genome reduction following the transition from nutrient-poor (groundwater or soil) to nutrient-rich (gut) environments. Genome reduction following formation of a long-term association with another organism has been well documented for many endosymbionts (for example of insects; [101] or in lineage I; [12]). However, for *Elusimicrobia*, genome reduction of free-living bacteria may have involved genome streamlining rather than genetic drift enabled by small population sizes in the intracellular environment.

Perhaps the most remarkable finding is the presence of a novel group of nitrogenase paralogs that are phylogenetically distinct from the five known groups. The new paralogs branch next to the group IV and V and co-occur with an extensive suite of radical SAM-based proteins in groundwater-associated *Elusimicrobia* from lineages IV and V. Based on gene context and protein phylogeny, it is unlikely the new paralog performs nitrogen fixation. Instead, we predict that the nifH proteins reduce a tetrapyrrole. Given the functions of related clades of genes, the nifH-like protein may synthesize a novel cofactor, but further investigations are needed to determine the product and its function. We anticipate exciting and perhaps unprecedented chemistry associated with this enzyme, given the versatility of nitrogenases and nitrogenase homologs in substrate reduction [75].

## Acknowledgements

Support was provided by grants from the Lawrence Berkeley National Laboratory’s Genomes-to-Watershed Scientific Focus Area. The U.S. Department of Energy (DOE), Office of Science, and Office of Biological and Environmental Research funded the work under contract DE-AC02-05CH11231 and the DOE carbon cycling program DOE-SC10010566, the Innovative Genomics Institute at Berkeley and the Chan Zuckerberg Biohub. The Ministry of Economy, Trade and Industry of Japan funded a part of the work as “The project for validating assessment methodology in geological disposal system”. Teruki Iwatsuki, Kazuki Hayashida, Toshihiro Kato, Mitsuru Kubota, Kazuya Miyakawa, and Akihito Mochizuki assisted with groundwater sampling at Mizunami and Horonobe Underground Research Laboratories, Japan Atomic Energy Agency (JAEA). C.H. acknowledges the Camille and Henry Dreyfus Foundation Postdoctoral Program in Environmental Chemistry for a Fellowship. L.A.H is supported by a Tier II Canada Research Chair.

## Conflict of interests

J.F.B. is a founder of Metagenomi. The other authors declare no competing interests.

## Author contributions

J.F.B., L.A.H and R.M. conceived the study. C.J.C., P.M.C., L.A.H. and R.M. analysed the genomic data. R.M. and L.A.H. performed the phylogenetic analyses. I.F. performed the Cazy analysis. L-X.C., Y.A., C.H., I.F. and J.F.B. provided the new genomes. R.M., J.F.B, C.J.C and P.M.C. wrote the manuscript with input from all authors. All documents were edited and approved by all authors.

## Data and Materials availability

The genomes of the herein analysed *Elusimicrobia* have been made publicly available on NCBI and on the ggkbase database (https://ggkbase.berkeley.edu/non_redundant_elusimicrobia_2018/organisms). Detailed annotations of the metabolic repertoire are provided in **Suppl. Dataset 3** accompanying this paper. Raw data files (phylogenetic trees, fasta sequences and the list of KEGG annotations) are made available via figshare under the following link: https://figshare.com/articles/Elusimicrobia_analysis/8939678. Correspondence and material requests should be addressed to.

## Notes

### Summary of Updates

we updated the title and the conflicts of interest

