## SupplementaryMaterials for "Groundwater *Elusimicrobia* are metabolically diverse compared to gut microbiome *Elusimicrobia* and some have a novel nitrogenase paralog"

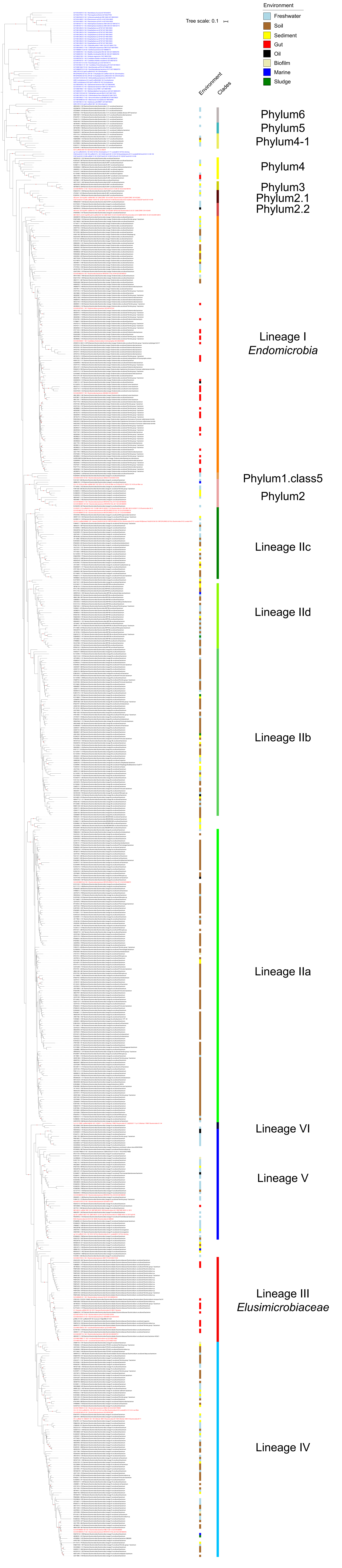

Environment
Freshwater
Soil
Sediment
Gut
Oil
Biofilm
Marine
Sludge
Tree scale: 0.1
Environment
Clades
Phylum6
Phylum5
Phylum4-1
Phylum4
Phylum3
Phylum2.1
Phylum2.2
Lineage I
Endomicrobia
Phylum1.class5
Phylum2
Lineage IIc
Lineage IId
Lineage IIb
Lineage IIa
Lineage VI
Lineage V
Lineage III
Elusimicrobiaceae
Lineage IV

Figure S1. Phylogenetic classification of 41 *Elusimicrobia*-related genomes based on their 16S rRNA gene sequences. Reference sequences from the SILVA database anchor the tree. The maximum likelihood phylogeny was inferred under the GTR model of evolution. For each *Elusimicrobia* genome, we indicated the lineage where the 16S sequence is nested in based on Herlemann *et al.* 2007; Geissinger *et al.* 2009 and Yarza *et al.* 2014. Scale bar indicates the average number of substitutions per site. The inner color bar indicates the environment of origin. Operational Taxonomic Unit (OTU) names colored in blue correspond to 16S from genome outgroups used in this study whereas OTUs colored in red correspond to the 16S from the Elusimicrobia genomes used in this study. OTUs colored in black were downloaded from the SILVA database.

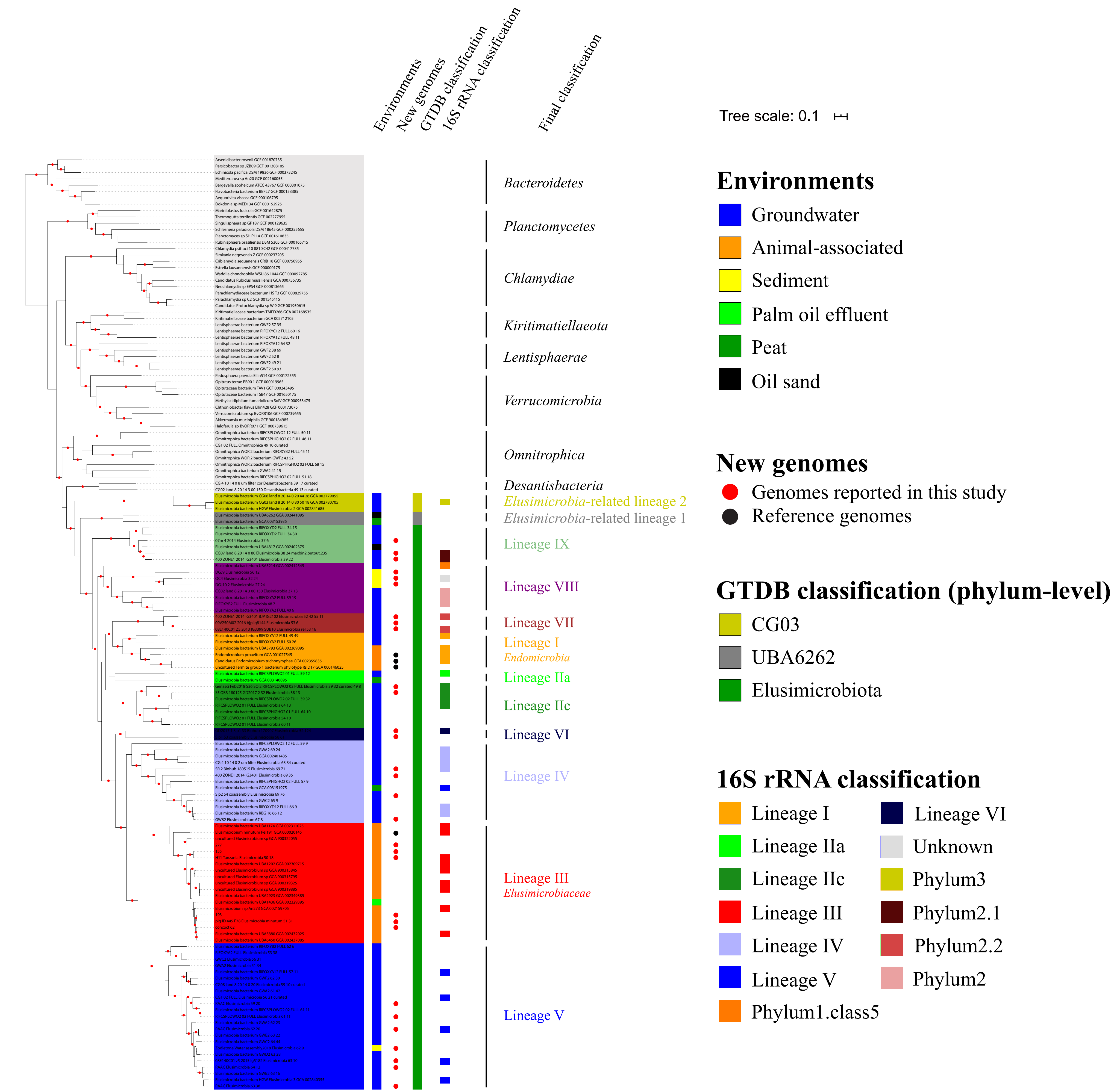

Environments
New genomes
GTDB classification
16S rRNA classification
Final classification
Tree scale: 0.1
Bacteroidetes
Planctomycetes
Chlamydiae
Kiritimatiellaeota
Lentisphaerae
Verrucomicrobia
Omnitrophica
Desantisbacteria
Elusimicrobia-related lineage 2
Elusimicrobia-related lineage 1
Lineage IX
Lineage VIII
Lineage VII
Lineage I
Endomicrobia
Lineage IIa
Lineage IIc
Lineage VI
Lineage IV
Lineage III
Elusimicrobiaceae
Lineage V
Environments
Groundwater
Animal-associated
Sediment
Palm oil effluent
Peat
Oil sand
New genomes
Genomes reported in this study
Reference genomes
GTDB classification (phylum-level)
CG03
UBA6262
Elusimicrobiota
16S rRNA classification
Lineage I
Lineage VI
Lineage IIa
Unknown
Lineage IIc
Phylum3
Lineage III
Phylum2.1
Lineage IV
Phylum2.2
Lineage V
Phylum2
Phylum1.class5

Figure S2. Placement of the 94 genomes related to the *Elusimicrobia* phylum. The maximum likelihood tree was constructed based on a concatenated alignment of 16 ribosomal proteins under an LG+gamma model of evolution. The inner color bar indicates the environment of origin. Black circles represent the four genomes described in previous studies whereas red circles represent the 30 new genomes added in this study. The remaining genomes come from the NCBI genome database. The two classifications based on 16S rRNA sequences and the GTDB are also indicated. Scale bars indicate the mean number of substitutions per site. The lineages of genomes are indicated by colored ranges and roman numbers. The bootstraps are indicated by red circles when ≥ 85.

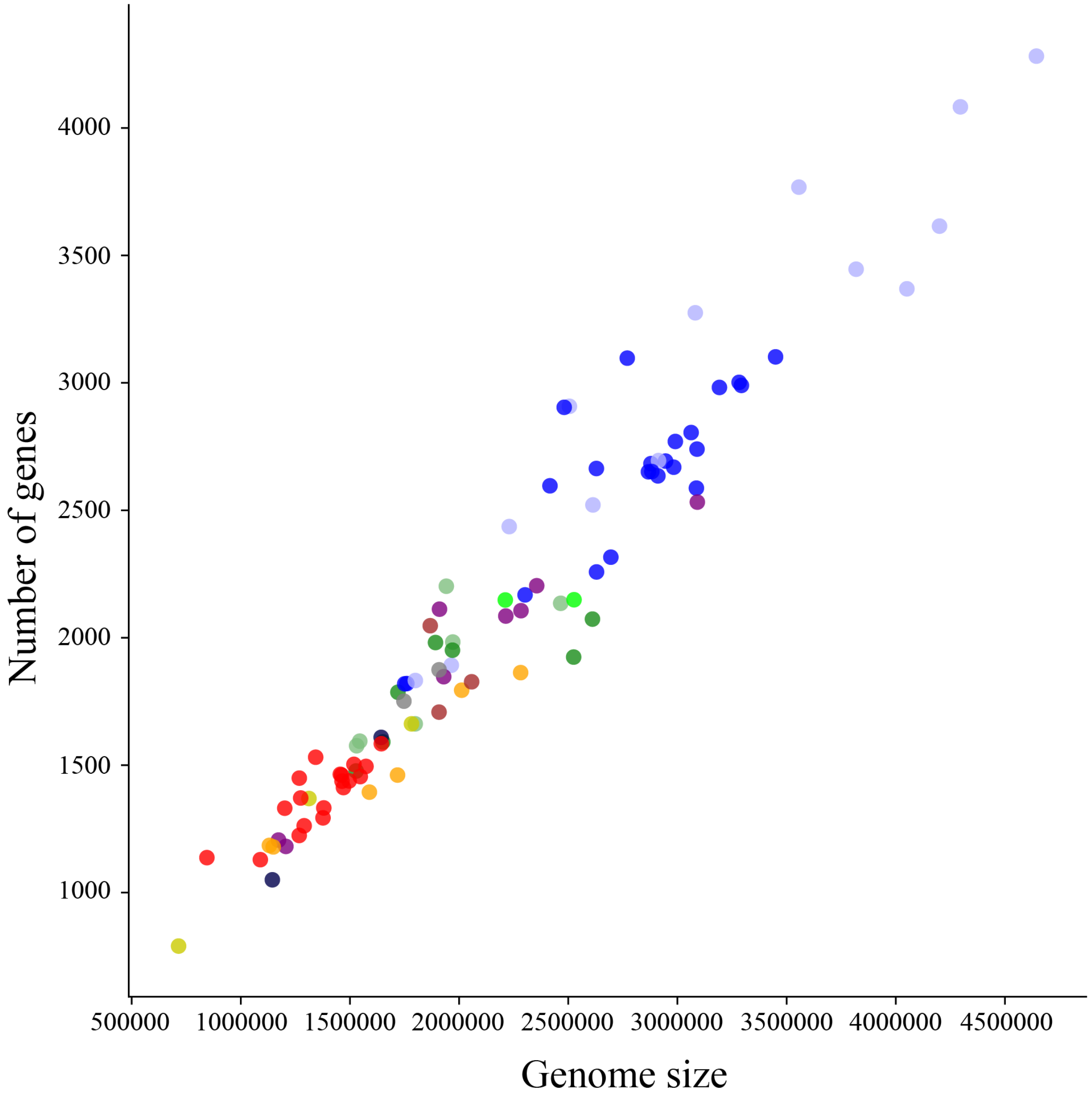


Figure S3. Number of genes and size (in base pairs) of the 94 *Elusimicrobia* genomes. Each genome is represented by a dot of the color of the lineage it has been classified to (see Figure 1 for color scheme).

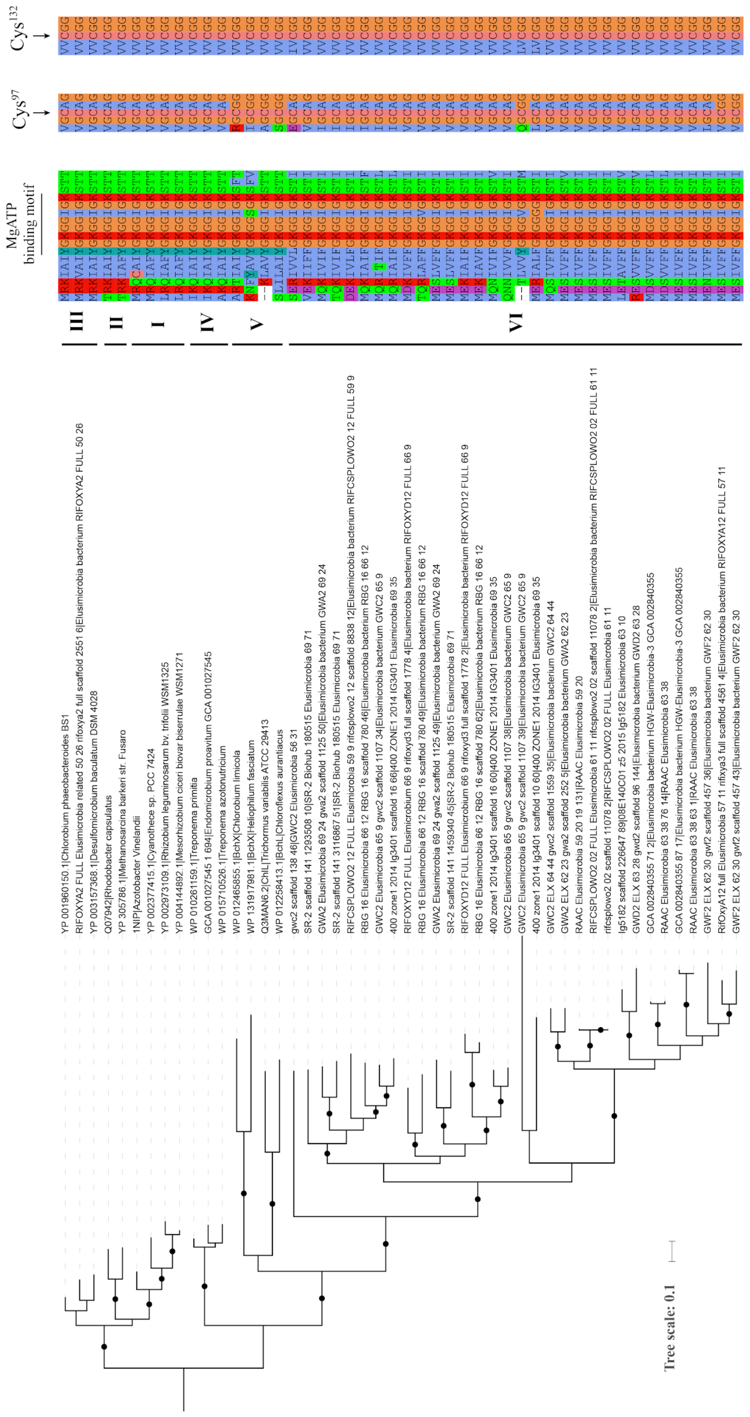


Figure S4. Phylogenetic tree of the nifH genes of *Elusimicrobia* and the nitrogenases of other bacterial species, and the conserved amino acids around the two crucial cysteine residues that coordinate the [$Fe_4S_4$] ligands in the Fe protein and for the binding of MgATP (GXGXXG). The cysteine residues, based on *A. vinelandii* numbering, are indicated with vertical arrows. Scale bar indicates substitutions per site.

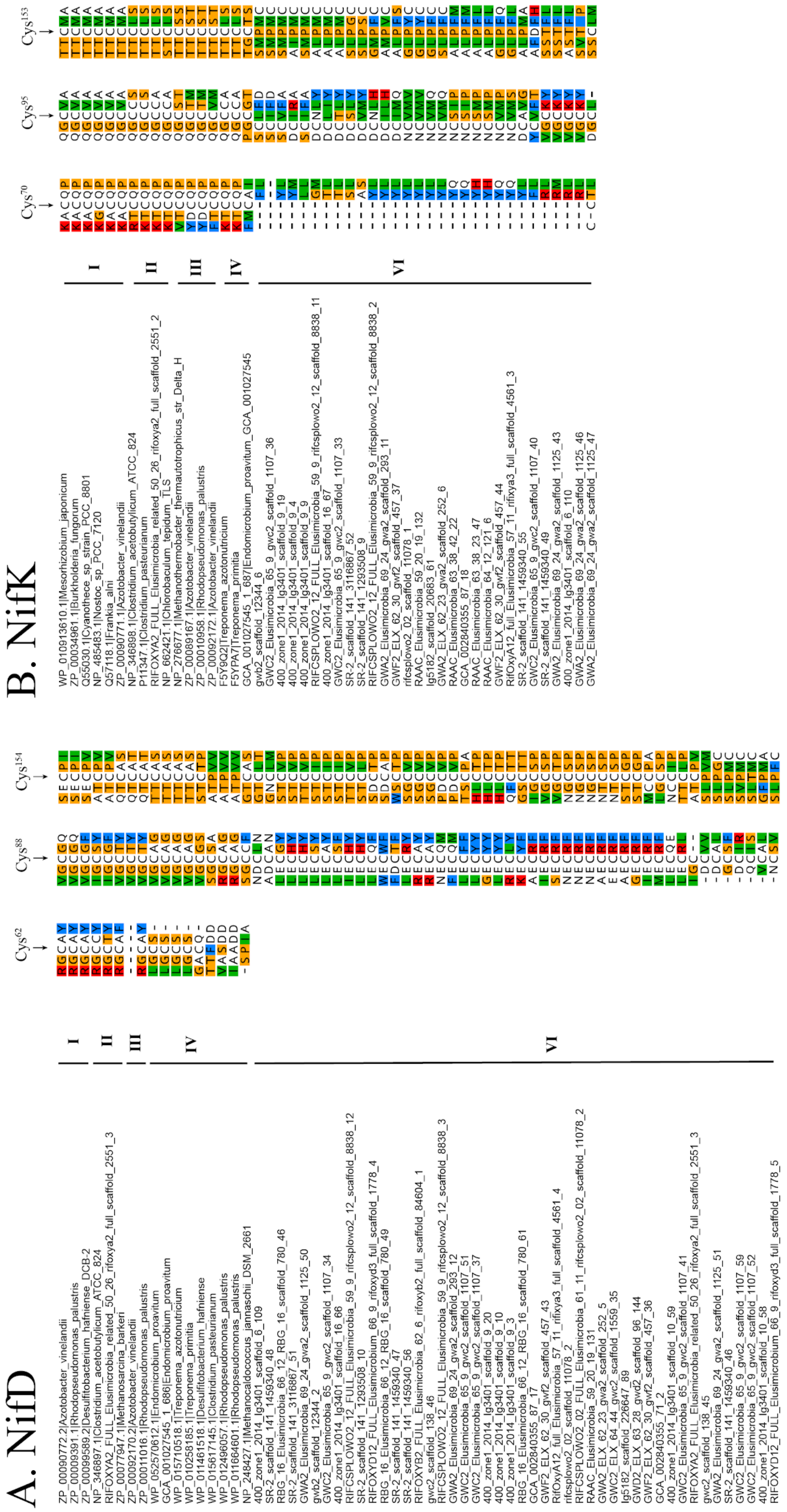


Figure S5. Conservation of the three crucial cysteine residues that coordinate the P-cluster in NifD (alpha) and NifK (beta) in *Elusimicrobia* and homologs in groups I, II, III and VI. The cysteine residues, based on *A vinelandii* numbering, are indicated with vertical arrows.

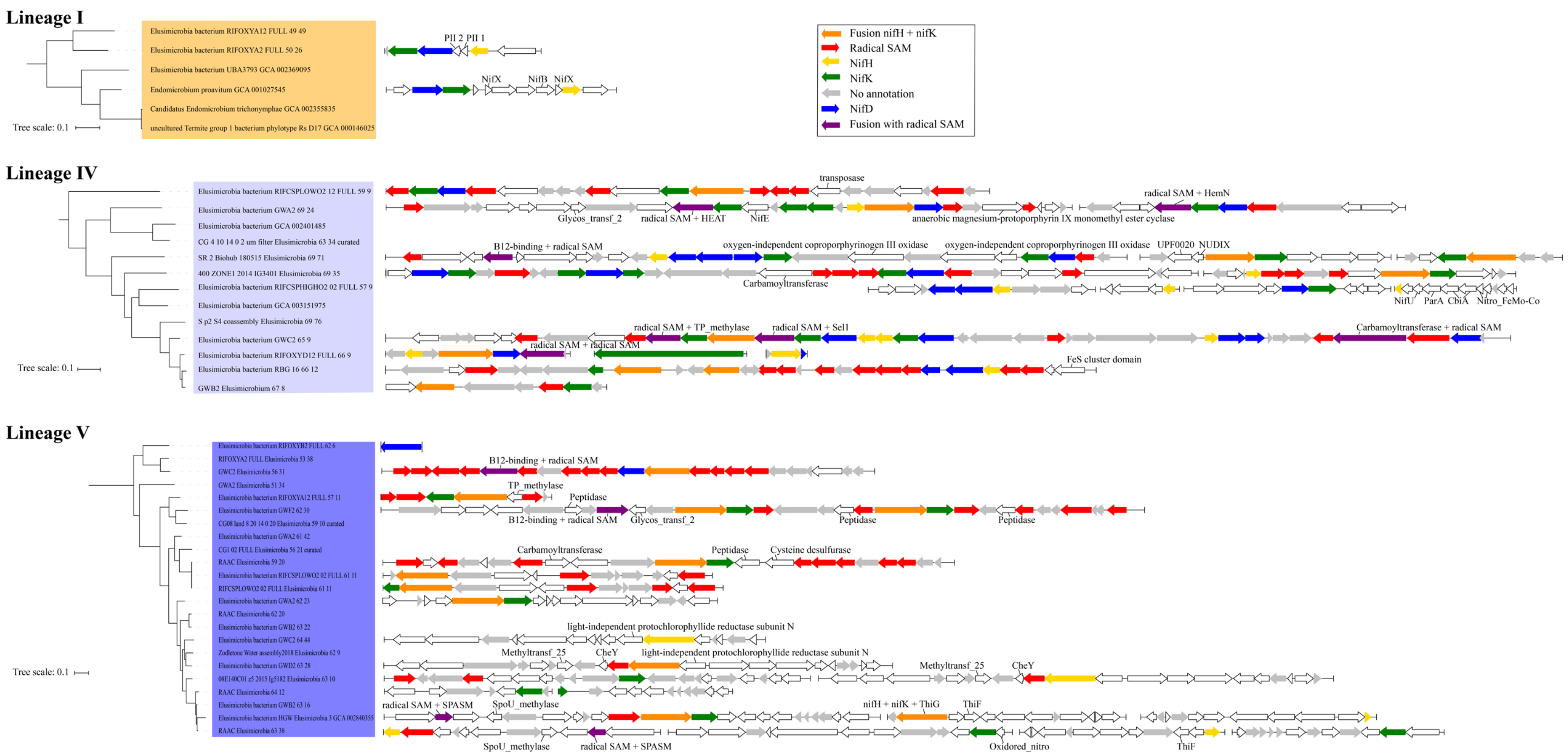
Lineage I
Elusimicrobia bacterium RIFOXYA12 FULL 49 49
Elusimicrobia bacterium RIFOXYA2 FULL 50 26
Elusimicrobia bacterium UBA3793 GCA 002369095
Endomicrobium proavitum GCA 001027545
Candidatus Endomicrobium trichonymphae GCA 002355835
uncultured Termite group 1 bacterium phylotype Rs D17 GCA 000146025
Tree scale: 0.1
PII 2 PII 1
NifX
NifB
NifX
Fusion nifH + nifK
Radical SAM
NifH
NifK
No annotation
NifD
Fusion with radical SAM
Lineage IV
Elusimicrobia bacterium RIFCSPLOWO2 12 FULL 59 9
Elusimicrobia bacterium GWA2 69 24
Elusimicrobia bacterium GCA 002401485
CG 4 10 14 0 2 um filter Elusimicrobia 63 34 curated
SR 2 Biohub 180515 Elusimicrobia 69 71
400 ZONE1 2014 IG3401 Elusimicrobia 69 35
Elusimicrobia bacterium RIFCSPHIGHO2 02 FULL 57 9
Elusimicrobia bacterium GCA 003151975
S p2 S4 coassembly Elusimicrobia 69 76
Elusimicrobia bacterium GWC2 65 9
Elusimicrobia bacterium RIFOXYD12 FULL 66 9
Elusimicrobia bacterium RBG 16 66 12
GWB2 Elusimicrobium 67 8
Tree scale: 0.1
transposase
Glycos_transf_2
radical SAM + HEAT
NifE
anaerobic magnesium-protoporphyrin IX monomethyl ester cyclase
radical SAM + HemN
B12-binding + radical SAM
oxygen-independent coproporphyrinogen III oxidase
oxygen-independent coproporphyrinogen III oxidase
UPF0020
NUDIX
Carbamoyltransferase
NifU
ParA
CbiA
Nitro_FeMo-Co
radical SAM + TP_methylase
radical SAM + Sell
Carbamoyltransferase + radical SAM
radical SAM + radical SAM
FeS cluster domain
Lineage V
Elusimicrobia bacterium RIFOXYB2 FULL 62 6
RIFOXYA2 FULL Elusimicrobia 53 38
GWC2 Elusimicrobia 56 31
GWA2 Elusimicrobia 51 34
Elusimicrobia bacterium RIFOXYA12 FULL 57 11
Elusimicrobia bacterium GWF2 62 30
CG08 land 8 20 14 0 20 Elusimicrobia 59 10 curated
Elusimicrobia bacterium GWA2 61 42
CG1 02 FULL Elusimicrobia 56 21 curated
RAAC Elusimicrobia 59 20
Elusimicrobia bacterium RIFCSPLOWO2 02 FULL 61 11
RIFCSPLOWO2 02 FULL Elusimicrobia 61 11
Elusimicrobia bacterium GWA2 62 23
RAAC Elusimicrobia 62 20
Elusimicrobia bacterium GWB2 63 22
Elusimicrobia bacterium GWC2 64 44
Zodletone Water assembly2018 Elusimicrobia 62 9
Elusimicrobia bacterium GWD2 63 28
08E140C01 z5 2015 lg5182 Elusimicrobia 63 10
RAAC Elusimicrobia 64 12
Elusimicrobia bacterium GWB2 63 16
Elusimicrobia bacterium HGW Elusimicrobia 3 GCA 002840355
RAAC Elusimicrobia 63 38
Tree scale: 0.1
B12-binding + radical SAM
TP_methylase
Peptidase
B12-binding + radical SAM
Glycos_transf_2
Peptidase
Peptidase
Carbamoyltransferase
Peptidase
Cysteine desulfurase
light-independent protochlorophyllide reductase subunit N
Methyltransf_25
CheY
light-independent protochlorophyllide reductase subunit N
Methyltransf_25
CheY
radical SAM + SPASM
SpoU_methylase
nifH + nifK + ThiG
ThiF
SpoU_methylase
radical SAM + SPASM
Oxidored_nitro
ThiF

Figure S6. Genomic neighborhood of nitrogenase in genomes from lineages I, IV and V. The 20 genes before and after the *nifH* genes are displayed when possible. Genes are colored according to their annotations. The maximum likelihood tree was constructed based on a concatenated alignment of 16 ribosomal proteins under an LG+gamma model of evolution (Figure 1). Scale bars indicate substitutions per site.

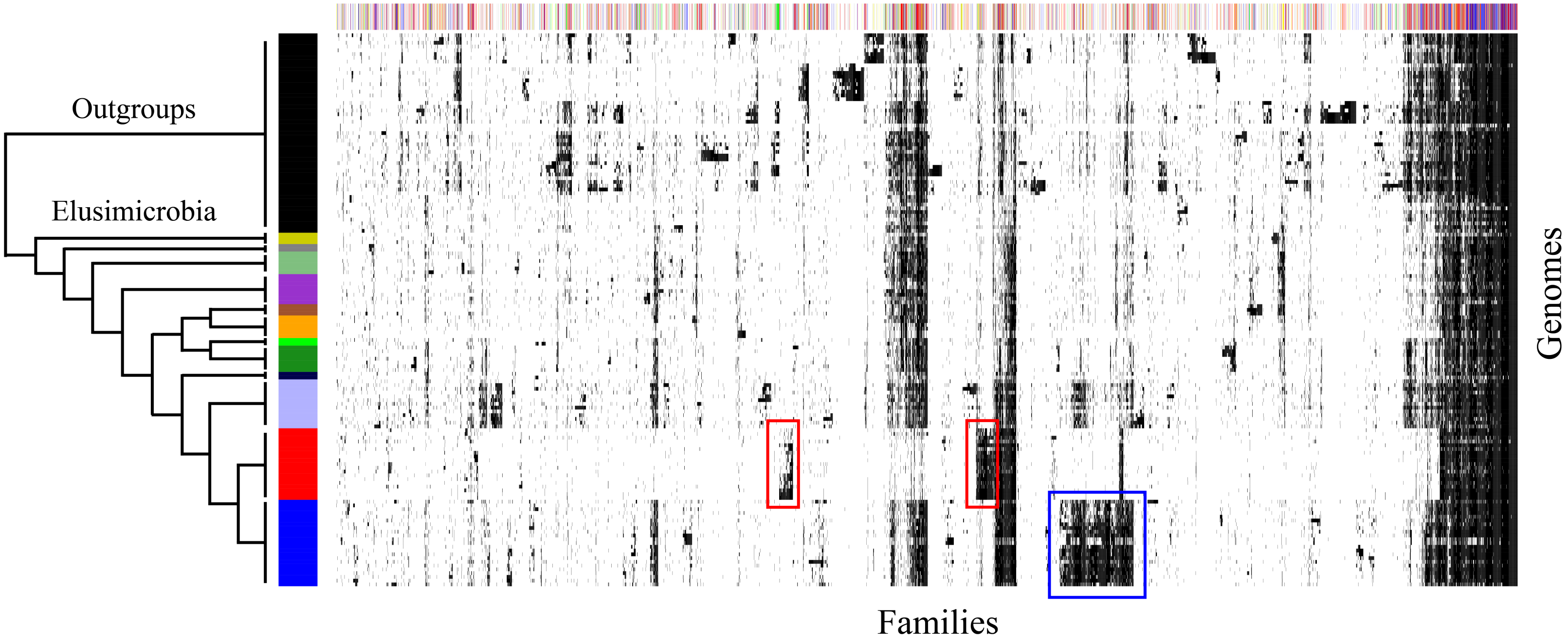
Outgroups
Elusimicrobia
Families
Genomes

'

Figure S7. The distribution of 6,608 protein families across the 110 genomes. The distribution of 6,608 protein families (columns) in 110 genomes (rows). Genomes were sorted according to the phylogeny in Figure 1. Families are clustered based on the presence (black) / absence (white) profiles (using Jaccard distance and complete-linkage method). The colored top bar corresponds to the functional category of families (Metabolism: red, Genetic Information Processing: blue, Cellular Processes: green, Environmental Information Processing: yellow, Organismal systems: orange, Unclassified: grey, Unknown: white). Specific clusters of interest are highlighted with colored boxes, discussed in the main text.

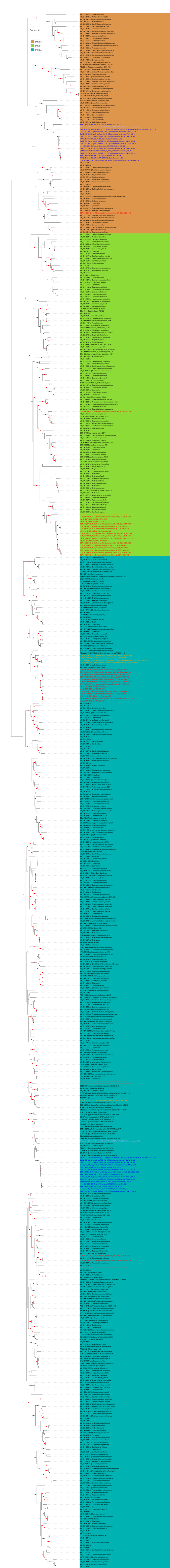

Tree scale: 0.1
group C
group B
group A

Figure S8. Phylogenetic tree of the [FeFe] hydrogenase (Maximum likelihood tree under an LG+I+G4 model of evolution). *Elusimicrobia* have group A, B and C [FeFe] hydrogenases. *Elusimicrobia* sequences are indicated with colored font. Scale bar indicates substitutions per site. The bootstraps are indicated by red circles when ≥ 85. The tree was made using sequences from (Greening et al. 2016; Matheus Carnevali et al. 2019). The sequences and tree are available with full bootstrap values in Fasta and Newick formats in Supplementary Data.

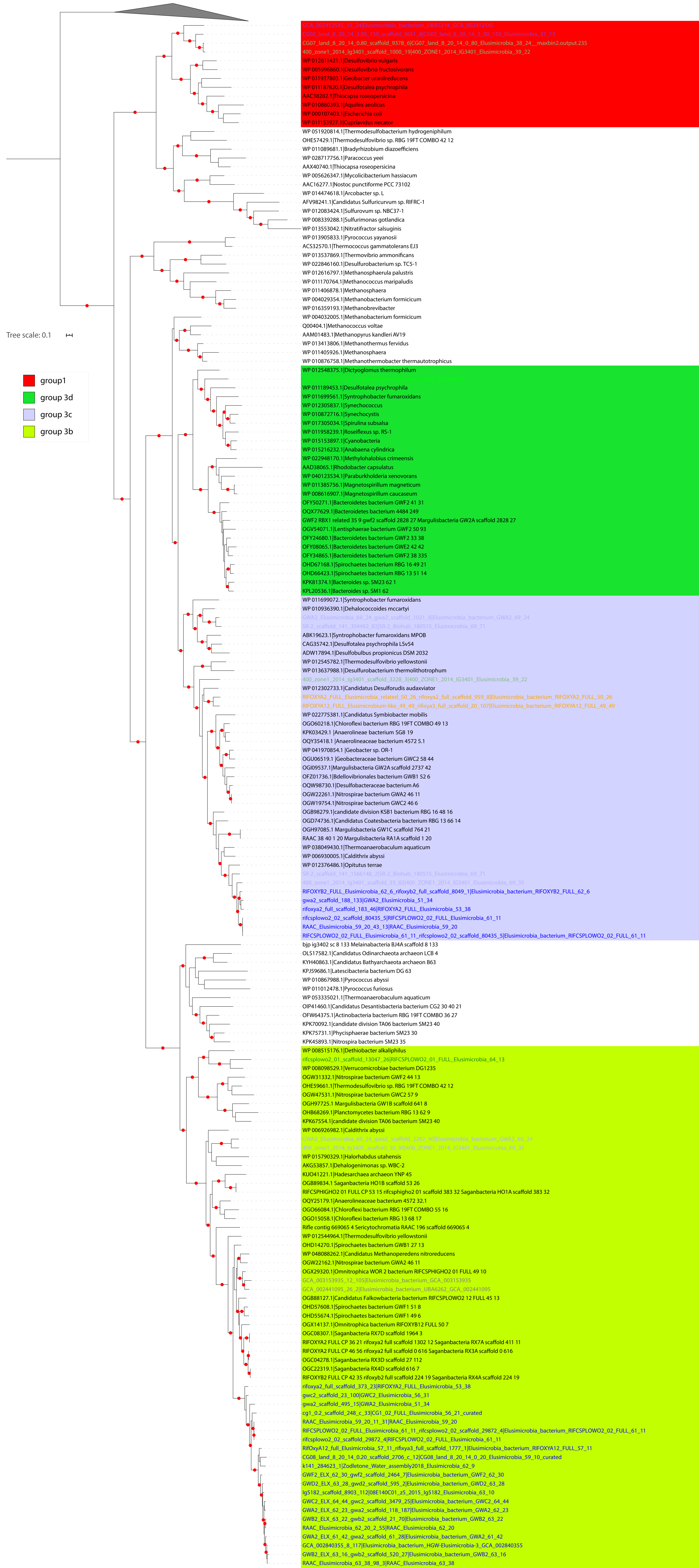

Tree scale: 0.1
group1
group 3d
group 3c
group 3b
GCA_002412545_15_24|Elusimicrobia_bacterium_UBA5214_GCA_002412545
CG02_land_8_20_14_3.00_150_scaffold_3651_8|CG02_land_8_20_14_3_00_150_Elusimicrobia_37_13
CG07_land_8_20_14_0.80_scaffold_9378_6|CG07_land_8_20_14_0_80_Elusimicrobia_38_24__maxbin2.output.235
400_zone1_2014_lg3401_scaffold_1000_19|400_ZONE1_2014_IG3401_Elusimicrobia_39_22
WP 012611421.1|Desulfovibrio vulgaris
WP 005996860.1|Desulfovibrio fructosivorans
WP 011937803.1|Geobacter uraniireducens
WP 011187820.1|Desulfotalea psychrophila
AAC38282.1|Thiocapsa roseopersicina
WP 010880393.1|Aquifex aeolicus
WP 000107403.1|Escherichia coli
WP 011153927.1|Cupriavidus necator
WP 051920814.1|Thermodesulfobacterium hydrogeniphilum
OHE57429.1|Thermodesulfovibrio sp. RBG 19FT COMBO 42 12
WP 011089681.1|Bradyrhizobium diazoefficiens
WP 028717756.1|Paracoccus yeei
AAX40740.1|Thiocapsa roseopersicina
WP 005626347.1|Mycolicibacterium hassiacum
AAC16277.1|Nostoc punctiforme PCC 73102
WP 014474618.1|Arcobacter sp. L
AFV98241.1|Candidatus Sulfuricurvum sp. RIFRC-1
WP 012083424.1|Sulfurovum sp. NBC37-1
WP 008339288.1|Sulfurimonas gotlandica
WP 013553042.1|Nitratifractor salsuginis
WP 013905833.1|Pyrococcus yayanosii
ACS32570.1|Thermococcus gammatolerans EJ3
WP 013537869.1|Thermovibrio ammonificans
WP 022846160.1|Desulfurobacterium sp. TC5-1
WP 012616797.1|Methanosphaerula palustris
WP 011170764.1|Methanococcus maripaludis
WP 011406878.1|Methanosphaera
WP 004029354.1|Methanobacterium formicicum
WP 016359193.1|Methanobrevibacter
WP 004032005.1|Methanobacterium formicicum
Q00404.1|Methanococcus voltae
AAM01483.1|Methanopyrus kandleri AV19
WP 013413806.1|Methanothermus fervidus
WP 011405926.1|Methanosphaera
WP 010876758.1|Methanothermobacter thermautotrophicus
WP 012548375.1|Dictyoglomus thermophilum
GCA_003140895_92_4|Elusimicrobia_bacterium_GCA_003140895
WP 011189453.1|Desulfotalea psychrophila
WP 011699561.1|Syntrophobacter fumaroxidans
WP 012305837.1|Synechococcus
WP 010872716.1|Synechocystis
WP 017305034.1|Spirulina subsalsa
WP 011958239.1|Roseiflexus sp. RS-1
WP 015153897.1|Cyanobacteria
WP 015216232.1|Anabaena cylindrica
WP 022948170.1|Methylohalobius crimeensis
AAD38065.1|Rhodobacter capsulatus
WP 040123534.1|Paraburkholderia xenovorans
WP 011385756.1|Magnetospirillum magneticum
WP 008616907.1|Magnetospirillum caucaseum
OFY50271.1|Bacteroidetes bacterium GWF2 41 31
OQX77629.1|Bacteroidetes bacterium 4484 249
GWF2 RBX1 related 35 9 gwf2 scaffold 2828 27 Margulisbacteria GW2A scaffold 2828 27
OGV54071.1|Lentisphaerae bacterium GWF2 50 93
OFY24680.1|Bacteroidetes bacterium GWF2 33 38
OFY08065.1|Bacteroidetes bacterium GWE2 42 42
OFY34865.1|Bacteroidetes bacterium GWF2 38 335
OHD67168.1|Spirochaetes bacterium RBG 16 49 21
OHD66423.1|Spirochaetes bacterium RBG 13 51 14
KPK81374.1|Bacteroides sp. SM23 62 1
KPL20536.1|Bacteroides sp. SM1 62
WP 011699072.1|Syntrophobacter fumaroxidans
WP 010936390.1|Dehalococcoides mccartyi
GWA2_Elusimicrobia_69_24_gwa2_scaffold_1021_8|Elusimicrobia_bacterium_GWA2_69_24
SR-2_scaffold_141_304462_82|SR-2_Biohub_180515_Elusimicrobia_69_71
ABK19623.1|Syntrophobacter fumaroxidans MPOB
CAG35742.1|Desulfotalea psychrophila LSv54
ADW17894.1|Desulfobulbus propionicus DSM 2032
WP 012545782.1|Thermodesulfovibrio yellowstonii
WP 013637988.1|Desulfurobacterium thermolithotrophum
400_zone1_2014_lg3401_scaffold_3228_3|400_ZONE1_2014_IG3401_Elusimicrobia_39_22
WP 012302733.1|Candidatus Desulforudis audaxviator
RIFOXYA2_FULL_Elusimicrobia_related_50_26_rifoxya2_full_scaffold_959_8|Elusimicrobia_bacterium_RIFOXYA2_FULL_50_26
RIFOXYA12_FULL_Elusimicrobium-like_49_49_rifixya3_full_scaffold_20_107|Elusimicrobia_bacterium_RIFOXYA12_FULL_49_49
WP 022775381.1|Candidatus Symbiobacter mobilis
OGO60218.1|Chloroflexi bacterium RBG 19FT COMBO 49 13
KPK03429.1 |Anaerolineae bacterium SG8 19
OQY35418.1 |Anaerolineaceae bacterium 4572 5.1
WP 041970854.1 |Geobacter sp. OR-1
OGU06519.1 |Geobacteraceae bacterium GWC2 58 44
OGI09537.1|Margulisbacteria GW2A scaffold 2737 42
OFZ01736.1|Bdellovibrionales bacterium GWB1 52 6
OQW98730.1|Desulfobacteraceae bacterium A6
OGW22261.1|Nitrospirae bacterium GWA2 46 11
OGW19754.1|Nitrospirae bacterium GWC2 46 6
OGB98279.1|candidate division KSB1 bacterium RBG 16 48 16
OGD74736.1|Candidatus Coatesbacteria bacterium RBG 13 66 14
OGH97085.1 Margulisbacteria GW1C scaffold 764 21
RAAC 38 40 1 20 Margulisbacteria RA1A scaffold 1 20
WP 038049430.1|Thermoanaerobaculum aquaticum
WP 006930005.1|Caldithrix abyssi
WP 012376486.1|Opitutus terrae
SR-2_scaffold_141_1566148_2|SR-2_Biohub_180515_Elusimicrobia_69_71
400_zone1_2014_lg3401_scaffold_33_83|400_ZONE1_2014_IG3401_Elusimicrobia_69_35
RIFOXYB2_FULL_Elusimicrobia_62_6_rifoxyb2_full_scaffold_8049_1|Elusimicrobia_bacterium_RIFOXYB2_FULL_62_6
gwa2_scaffold_188_133|GWA2_Elusimicrobia_51_34
rifoxya2_full_scaffold_183_46|RIFOXYA2_FULL_Elusimicrobia_53_38
rifcsplowo2_02_scaffold_80435_5|RIFCSPLOWO2_02_FULL_Elusimicrobia_61_11
RAAC_Elusimicrobia_59_20_43_13|RAAC_Elusimicrobia_59_20
RIFCSPLOWO2_02_FULL_Elusimicrobia_61_11_rifcsplowo2_02_scaffold_80435_5|Elusimicrobia_bacterium_RIFCSPLOWO2_02_FULL_61_11
bjp ig3402 sc 8 133 Melainabacteria BJ4A scaffold 8 133
OLS17582.1|Candidatus Odinarchaeota archaeon LCB 4
KYH40863.1|Candidatus Bathyarchaeota archaeon B63
KPJ59686.1|Latescibacteria bacterium DG 63
WP 010867988.1|Pyrococcus abyssi
WP 011012478.1|Pyrococcus furiosus
WP 053335021.1|Thermoanaerobaculum aquaticum
OIP41460.1|Candidatus Desantisbacteria bacterium CG2 30 40 21
OFW64375.1|Actinobacteria bacterium RBG 19FT COMBO 36 27
KPK70092.1|candidate division TA06 bacterium SM23 40
KPK75731.1|Phycisphaerae bacterium SM23 30
KPK45893.1|Nitrospira bacterium SM23 35
WP 008515176.1|Dethiobacter alkaliphilus
rifcsplowo2_01_scaffold_13047_26|RIFCSPLOWO2_01_FULL_Elusimicrobia_64_13
WP 008098529.1|Verrucomicrobiae bacterium DG1235
OGW31332.1|Nitrospirae bacterium GWF2 44 13
OHE59661.1|Thermodesulfovibrio sp. RBG 19FT COMBO 42 12
OGW47531.1|Nitrospirae bacterium GWC2 57 9
OGH97725.1 Margulisbacteria GW1B scaffold 641 8
OHB68269.1|Planctomycetes bacterium RBG 13 62 9
KPK67554.1|candidate division TA06 bacterium SM23 40
WP 006926982.1|Caldithrix abyssi
GWA2_Elusimicrobia_69_24_gwa2_scaffold_2292_40|Elusimicrobia_bacterium_GWA2_69_24
400_zone1_2014_lg3401_scaffold_30_90|400_ZONE1_2014_IG3401_Elusimicrobia_69_35
WP 015790329.1|Halorhabdus utahensis
AKG53857.1|Dehalogenimonas sp. WBC-2
KUO41221.1|Hadesarchaea archaeon YNP 45
OGB89834.1 Saganbacteria HO1B scaffold 53 26
RIFCSPHIGHO2 01 FULL CP 53 15 rifcsphigho2 01 scaffold 383 32 Saganbacteria HO1A scaffold 383 32
OQY25179.1|Anaerolineaceae bacterium 4572 32.1
OGO66084.1|Chloroflexi bacterium RBG 19FT COMBO 55 16
OGO15058.1|Chloroflexi bacterium RBG 13 68 17
Rifle contig 669065 4 Sericytochromatia RAAC 196 scaffold 669065 4
WP 012544964.1|Thermodesulfovibrio yellowstonii
OHD14270.1|Spirochaetes bacterium GWB1 27 13
WP 048088262.1|Candidatus Methanoperedens nitroreducens
OGW22162.1|Nitrospirae bacterium GWA2 46 11
OGX29320.1|Omnitrophica WOR 2 bacterium RIFCSPHIGHO2 01 FULL 49 10
GCA_003153935_12_105|Elusimicrobia_bacterium_GCA_003153935
GCA_002441095_26_2|Elusimicrobia_bacterium_UBA6262_GCA_002441095
OGB88127.1|Candidatus Falkowbacteria bacterium RIFCSPLOWO2 12 FULL 45 13
OHD57608.1|Spirochaetes bacterium GWF1 51 8
OHD55674.1|Spirochaetes bacterium GWF1 49 6
OGX14137.1|Omnitrophica bacterium RIFOXYB12 FULL 50 7
OGC08307.1|Saganbacteria RX7D scaffold 1964 3
RIFOXYA2 FULL CP 36 21 rifoxya2 full scaffold 1302 12 Saganbacteria RX7A scaffold 411 11
RIFOXYA2 FULL CP 46 56 rifoxya2 full scaffold 0 616 Saganbacteria RX3A scaffold 0 616
OGC04278.1|Saganbacteria RX3D scaffold 27 112
OGC22319.1|Saganbacteria RX4D scaffold 616 7
RIFOXYB2 FULL CP 42 35 rifoxyb2 full scaffold 224 19 Saganbacteria RX4A scaffold 224 19
rifoxya2_full_scaffold_373_23|RIFOXYA2_FULL_Elusimicrobia_53_38
gwc2_scaffold_23_100|GWC2_Elusimicrobia_56_31
gwa2_scaffold_495_15|GWA2_Elusimicrobia_51_34
cg1_0.2_scaffold_248_c_33|CG1_02_FULL_Elusimicrobia_56_21_curated
RAAC_Elusimicrobia_59_20_11_31|RAAC_Elusimicrobia_59_20
RIFCSPLOWO2_02_FULL_Elusimicrobia_61_11_rifcsplowo2_02_scaffold_29872_4|Elusimicrobia_bacterium_RIFCSPLOWO2_02_FULL_61_11
rifcsplowo2_02_scaffold_29872_4|RIFCSPLOWO2_02_FULL_Elusimicrobia_61_11
RifOxyA12_full_Elusimicrobia_57_11_rifixya3_full_scaffold_1777_1|Elusimicrobia_bacterium_RIFOXYA12_FULL_57_11
CG08_land_8_20_14_0.20_scaffold_2706_c_12|CG08_land_8_20_14_0_20_Elusimicrobia_59_10_curated
k141_284623_1|Zodletone_Water_assembly2018_Elusimicrobia_62_9
GWF2_ELX_62_30_gwf2_scaffold_2464_7|Elusimicrobia_bacterium_GWF2_62_30
GWD2_ELX_63_28_gwd2_scaffold_595_2|Elusimicrobia_bacterium_GWD2_63_28
lg5182_scaffold_8903_112|08E140C01_z5_2015_lg5182_Elusimicrobia_63_10
GWC2_ELX_64_44_gwc2_scaffold_3479_25|Elusimicrobia_bacterium_GWC2_64_44
GWA2_ELX_62_23_gwa2_scaffold_118_187|Elusimicrobia_bacterium_GWA2_62_23
GWB2_ELX_63_22_gwb2_scaffold_21_70|Elusimicrobia_bacterium_GWB2_63_22
RAAC_Elusimicrobia_62_20_2_55|RAAC_Elusimicrobia_62_20
GWA2_ELX_61_42_gwa2_scaffold_61_28|Elusimicrobia_bacterium_GWA2_61_42
GCA_002840355_8_117|Elusimicrobia_bacterium_HGW-Elusimicrobia-3_GCA_002840355
GWB2_ELX_63_16_gwb2_scaffold_520_27|Elusimicrobia_bacterium_GWB2_63_16
RAAC_Elusimicrobia_63_38_98_3|RAAC_Elusimicrobia_63_38

Figure S9. Phylogenetic tree of the groups 1, 2 and 3 of [NiFe] hydrogenase (Maximum likelihood tree under a LG+G4 model of evolution). *Elusimicrobia* have group 1, group 3b, 3c and 3d [NiFe] hydrogenases. *Elusimicrobia* sequences are indicated with colored font. Scale bar indicates substitutions per site. The bootstraps are indicated by red circles when ≥ 85. The tree was made using sequences from (Matheus Carnevali et al. 2019). The sequences and tree are available with full bootstrap values in Fasta and Newick format in Supplementary Data.

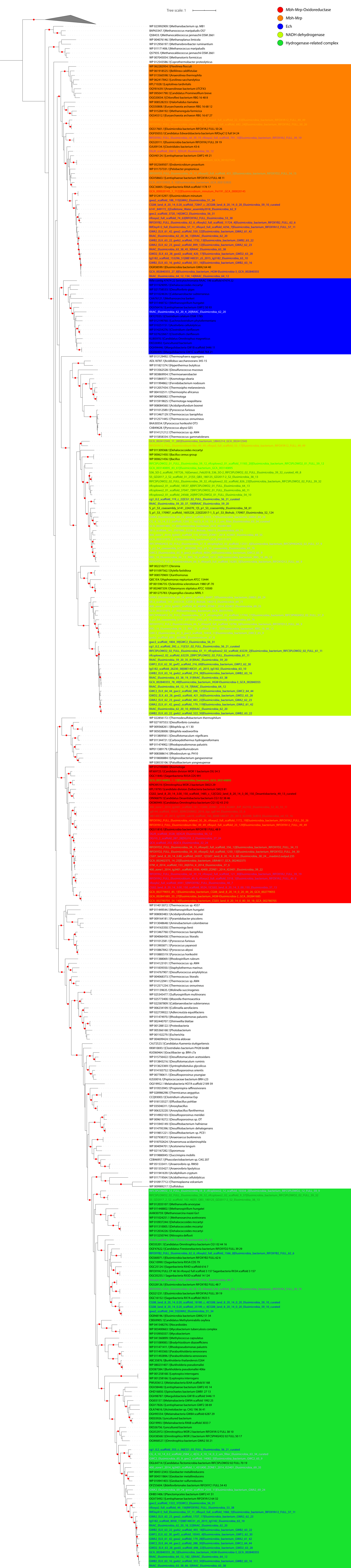
Tree scale: 1
Mbh-Mrp-Oxidoreductase
Mbh-Mrp
Ech
NADH dehydrogenase
Hydrogenase-related complex

Figure S10. Phylogenetic tree of the group 4 [NiFe] hydrogenase (Maximum likelihood tree under a LG+I+G4 model of evolution). *Elusimicrobia* have Mbh-Mrp, Mbh-Mrp-Oxidoreductase, Ech and Hydrogenase-related complex [NiFe] hydrogenases. They also have NADH dehydrogenase. *Elusimicrobia* sequences are indicated with colored font. Scale bar indicates substitutions per site. The bootstraps are indicated by red circles when ≥ 85. The tree was made using sequences from (Matheus Carnevali et al. 2019). The sequences and tree are available with full bootstrap values in Fasta and Newick format in Supplementary Data.

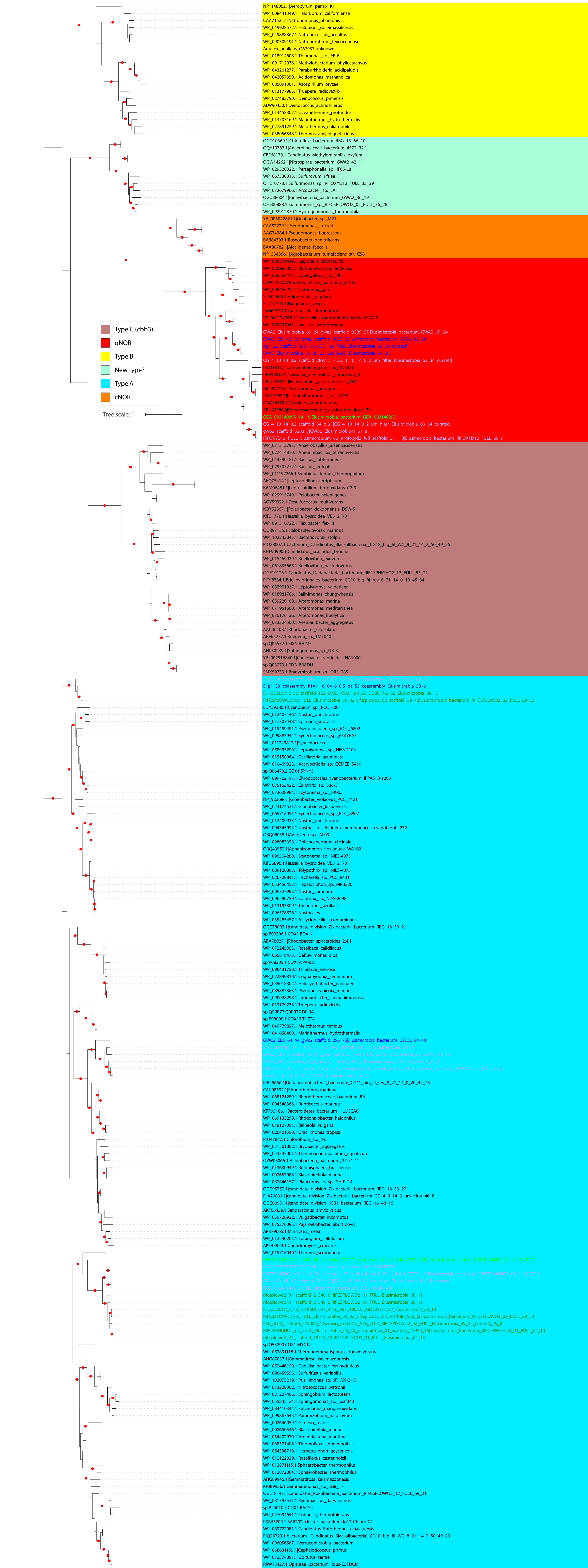

Type C (cbb3)
qNOR
Type B
New type?
Type A
cNOR
Tree scale: 1
NP_148062.1|Aeropyrum_pernix_K1
WP_008441349.1|Halorubrum_californiense
CAA71525.1|Natronomonas_pharaonis
WP_049928572.1|Halopiger_goleimassiliensis
WP_049888867.1|Natronococcus_occultus
WP_090309191.1|Natronorubrum_texcoconense
Aquifex_aeolicus_O67937|unknown
WP_018914608.1|Thiomonas_sp._FB-6
WP_091712936.1|Methylobacterium_phyllostachyos
WP_043201277.1|Paraburkholderia_acidipaludis
WP_042057359.1|Acidomonas_methanolica
WP_085091361.1|Azospirillum_oryzae
WP_013177985.1|Truepera_radiovictrix
WP_027483790.1|Deinococcus_pimensis
ALW90450.1|Deinococcus_actinosclerus
WP_013458307.1|Oceanithermus_profundus
WP_013703169.1|Marinithermus_hydrothermalis
WP_027891229.1|Meiothermus_chliarophilus
WP_038056548.1|Thermus_amyloliquefaciens
OGO10369.1|Chloroflexi_bacterium_RBG_13_66_10
OQY19783.1|Anaerolineaceae_bacterium_4572_32.1
CBE68178.1|Candidatus_Methylomirabilis_oxyfera
OGW14262.1|Nitrospirae_bacterium_GWA2_42_11
WP_029520322.1|Persephonella_sp._IF05-L8
WP_067330013.1|Sulfurovum_riftiae
OHE10778.1|Sulfurimonas_sp._RIFOXYD12_FULL_33_39
WP_072679966.1|Arcobacter_sp._LA11
OGU38604.1|Ignavibacteria_bacterium_GWA2_36_19
OHE00888.1|Sulfurimonas_sp._RIFCSPLOWO2_02_FULL_36_28
WP_092912870.1|Hydrogenimonas_thermophila
YP_003022821.1|Geobacter_sp._M21
CAA82229.1|Pseudomonas_stutzeri
AAG34384.1|Pseudomonas_fluorescens
BAB84303.1|Roseobacter_denitrificans
BAA90782.1|Alcaligenes_faecalis
NP_534866.1|Agrobacterium_tumefaciens_str._C58
GWA2_Elusimicrobia_69_24_gwa2_scaffold_3589_27|Elusimicrobia_bacterium_GWA2_69_24
CG_4_10_14_0.2_scaffold_2897_c_7|CG_4_10_14_0_2_um_filter_Elusimicrobia_63_34_curated
CG_4_10_14_0.2_scaffold_34_c_21|CG_4_10_14_0_2_um_filter_Elusimicrobia_63_34_curated
gwb2_scaffold_3283_7|GWB2_Elusimicrobium_67_8
RIFOXYD12_FULL_Elusimicrobium_66_9_rifoxyd3_full_scaffold_3111_3|Elusimicrobia_bacterium_RIFOXYD12_FULL_66_9
WP_071313791.1|Anaerobacillus_arsenicselenatis
WP_027414870.1|Aneurinibacillus_terranovensis
WP_044390181.1|Bacillus_subterraneus
WP_079507272.1|Bacillus_jeotgali
WP_011197266.1|Symbiobacterium_thermophilum
AEQ75414.1|Leptospirillum_ferriphilum
BAM06481.1|Leptospirillum_ferrooxidans_C2-3
WP_029915749.1|Pelobacter_selenigenes
AOY59322.1|Desulfococcus_multivorans
KOY52867.1|Polaribacter_dokdonensis_DSW-5
KIF31770.1|Hassallia_byssoidea_VB512170
WP_091516722.1|Flexibacter_flexilis
OUR97135.1|Halobacteriovorax_marinus
WP_102243045.1|Bacteriovorax_stolpii
PIQ28007.1|bacterium_(Candidatus_Blackallbacteria)_CG18_big_fil_WC_8_21_14_2_50_49_26
KHE90990.1|Candidatus_Scalindua_brodae
WP_015469929.1|Bdellovibrio_exovorus
WP_061835668.1|Bdellovibrio_bacteriovorus
OGE19126.1|Candidatus_Dadabacteria_bacterium_RIFCSPHIGHO2_12_FULL_53_21
PIT98784.1|Bdellovibrionales_bacterium_CG10_big_fil_rev_8_21_14_0_10_45_34
WP_082901917.1|Leptolyngbya_valderiana
WP_018981786.1|Salinimonas_chungwhensis
WP_039220109.1|Alteromonas_marina
WP_071951600.1|Alteromonas_mediterranea
WP_070176136.1|Alteromonas_lipolytica
WP_073324560.1|Aestuariibacter_aggregatus
AAC46108.1|Rhodobacter_capsulatus
ABF65277.1|Ruegeria_sp._TM1040
sp Q05572.1 FIXN RHIME
AHL30239.1|Sphingomonas_sp._NX-3
YP_002516840.1|Caulobacter_vibrioides_NA1000
sp Q03073.1 FIXN BRADU
SMX59739.1|Bradyrhizobium_sp._ORS_285
S_p1_S3_coassembly_k141_1616416_8|S_p1_S3_coassembly_Elusimicrobia_58_61
SS_GD2017_2_S2_scaffold_122_60|SS_QB3_180125_GD2017-2_S2_Elusimicrobia_38_13
RIFCSPLOWO2_02_FULL_Elusimicrobia_39_32_rifcsplowo2_02_scaffold_39_42|Elusimicrobia_bacterium_RIFCSPLOWO2_02_FULL_39_32
EDY39380.1|Cyanobium_sp._PCC_7001
WP_012407146.1|Nostoc_punctiforme
WP_017303448.1|Spirulina_subsalsa
WP_019499491.1|Pseudanabaena_sp._PCC_6802
WP_099883944.1|Synechococcus_sp._65AY6A5
WP_011243817.1|Synechococcus
WP_058995290.1|Leptolyngbya_sp._NIES-2104
WP_015150964.1|Oscillatoria_acuminata
WP_010469823.1|Acaryochloris_sp._CCMEE_5410
sp Q06473.2 COX1 SYNY3
WP_099702197.1|Chroococcales_cyanobacterium_IPPAS_B-1203
WP_035153422.1|Calothrix_sp._336/3
WP_073628984.1|Scytonema_sp._HK-05
NP_923686.1|Gloeobacter_violaceus_PCC_7421
WP_023174521.1|Gloeobacter_kilaueensis
WP_065716051.1|Synechococcus_sp._PCC_8807
WP_012409913.1|Nostoc_punctiforme
WP_094345093.1|Nostoc_sp._'Peltigera_membranacea_cyanobiont'_232
OBQ08035.1|Anabaena_sp._AL09
WP_028083358.1|Dolichospermum_circinale
OBQ43552.1|Aphanizomenon_flos-aquae_WA102
WP_096564280.1|Scytonema_sp._NIES-4073
KIF36896.1|Hassallia_byssoidea_VB512170
WP_089126893.1|Tolypothrix_sp._NIES-4075
WP_026720841.1|Fischerella_sp._PCC_9431
WP_053456453.1|Hapalosiphon_sp._MRB220
WP_096727993.1|Nostoc_carneum
WP_096590759.1|Calothrix_sp._NIES-2098
WP_013193309.1|Trichormus_azollae
WP_096578826.1|Nostocales
WP_035485457.1|Alicyclobacillus_contaminans
OGC78093.1|candidate_division_Zixibacteria_bacterium_RBG_16_50_21
sp P00396.1 COX1 BOVIN
ABA78027.1|Rhodobacter_sphaeroides_2.4.1
WP_072245353.1|Roseibaca_calidilacus
WP_066818473.1|Defluviimonas_alba
sp P08305.1 COX1A PARDE
WP_096431750.1|Thioclava_arenosa
WP_072899810.1|Cognatiyoonia_sediminum
WP_039018302.1|Halocynthiibacter_namhaensis
WP_085887363.1|Pseudooceanicola_marinus
WP_090028298.1|Lutimaribacter_saemankumensis
WP_013179238.1|Truepera_radiovictrix
sp Q9RR77 Q9RR77 DEIRA
sp P98005.1 COX13 THET8
WP_040779827.1|Meiothermus_timidus
WP_041658460.1|Marinithermus_hydrothermalis
GWC2_ELX_64_44_gwc2_scaffold_296_75|Elusimicrobia_bacterium_GWC2_64_44
PIR25656.1|Deltaproteobacteria_bacterium_CG11_big_fil_rev_8_21_14_0_20_42_23
CAC08532.1|Rhodothermus_marinus
WP_068121289.1|Rhodothermaceae_bacterium_RA
WP_094548366.1|Rubricoccus_marinus
KPP95186.1|Bacteroidetes_bacterium_HLUCCA01
WP_069133290.1|Rhodohalobacter_halophilus
WP_018127091.1|Balneola_vulgaris
WP_020401590.1|Gracilimonas_tropica
PIO47847.1|Chlorobium_sp._445
WP_031501002.1|Bryobacter_aggregatus
WP_053335001.1|Thermoanaerobaculum_aquaticum
OYW03066.1|Acidobacteria_bacterium_37-71-11
WP_013630949.1|Rubinisphaera_brasiliensis
WP_002653988.1|Blastopirellula_marina
WP_082846157.1|Planctomyces_sp._SH-PL14
OGC95752.1|candidate_division_Zixibacteria_bacterium_RBG_16_53_22
PJA26831.1|candidate_division_Zixibacteria_bacterium_CG_4_9_14_3_um_filter_46_8
OGC00951.1|candidate_division_KSB1_bacterium_RBG_16_48_16
AKF06459.1|Sandaracinus_amylolyticus
WP_050726922.1|Vulgatibacter_incomptus
WP_057276095.1|Pajaroellobacter_abortibovis
APR79665.1|Minicystis_rosea
WP_012240201.1|Sorangium_cellulosum
AKT43929.1|Chondromyces_crocatus
WP_015716040.1|Thermus_scotoductus
rifcsplowo2_01_scaffold_21348_6|RIFCSPLOWO2_01_FULL_Elusimicrobia_60_11
rifcsplowo2_01_scaffold_21348_5|RIFCSPLOWO2_01_FULL_Elusimicrobia_60_11
SS_GD2017_2_S2_scaffold_637_4|SS_QB3_180125_GD2017-2_S2_Elusimicrobia_38_13
RIFCSPLOWO2_02_FULL_Elusimicrobia_39_32_rifcsplowo2_02_scaffold_973_6|Elusimicrobia_bacterium_RIFCSPLOWO2_02_FULL_39_32
S36_SO-2_scaffold_379086_4|Genasci_Feb2018_S36_SO-2_RIFCSPLOWO2_02_FULL_Elusimicrobia_39_32_curated_49_8
RIFCSPHIGHO2_01_FULL_Elusimicrobia_64_10_rifcsphigho2_01_scaffold_19950_15|Elusimicrobia_bacterium_RIFCSPHIGHO2_01_FULL_64_10
rifcsplowo2_01_scaffold_19516_11|RIFCSPLOWO2_01_FULL_Elusimicrobia_64_13
sp Q53290 COX1 MYCTU
WP_052891118.1|Thermogemmatispora_carboxidivorans
AHG87837.1|Gemmatirosa_kalamazoonesis
WP_052446190.1|Geoalkalibacter_ferrihydriticus
WP_096459593.1|Sulfurifustis_variabilis
WP_102073219.1|Pusillimonas_sp._JR1/69-3-13
WP_013220362.1|Nitrosococcus_watsonii
WP_021227406.1|Sphingobium_lactosutens
WP_055845124.1|Sphingomonas_sp._Leaf343
WP_084410344.1|Fulvimarina_manganoxydans
WP_099867643.1|Parahizobium_haloflavum
WP_002646059.1|Gimesia_maris
WP_002650546.1|Blastopirellula_marina
WP_054492930.1|Ardenticatena_maritima
WP_088571408.1|Thermoflexus_hugenholtzii
WP_054536710.1|Herpetosiphon_geysericola
WP_012122029.1|Roseiflexus_castenholzii
WP_012871115.1|Sphaerobacter_thermophilus
WP_012872064.1|Sphaerobacter_thermophilus
AHG89945.1|Gemmatirosa_kalamazoonesis
KPJ89936.1|Gemmatimonas_sp._SG8_17
OGL18143.1|Candidatus_Rokubacteria_bacterium_RIFCSPLOWO2_12_FULL_69_21
WP_081793531.1|Paenibacillus_darwinianus
sp P24010.3 COX1 BACSU
WP_027094641.1|Cohnella_thermotolerans
PKB63209.1|SAR202_cluster_bacterium_Io17-Chloro-G1
WP_089722081.1|Candidatus_Entotheonella_palauensis
PIQ26723.1|bacterium_(Candidatus_Blackallbacteria)_CG18_big_fil_WC_8_21_14_2_50_49_26
WP_096056567.1|Verrucomicrobia_bacterium
WP_068631155.1|Cephaloticoccus_primus
WP_012374891.1|Opitutus_terrae
PAW73427.1|Opitutae_bacterium_Tous-C5TDCM

Figure S11. Phylogenetic tree of the catalytic subunit I of heme-copper oxygen reductases (Maximum likelihood tree under a LG+F+I+G4 model of evolution). *Elusimicrobia* have an $aa_3$-type heme-copper oxygen reductase (Type A) and nitric oxide reductase that uses quinol as the electron donor (qNOR). *Elusimicrobia* sequences are indicated with colored font. The bootstraps are indicated by red circles when ≥ 85. The tree was made using sequences from (Matheus Carnevali et al. 2019). Scale bar indicates substitutions per site. The sequences and tree are available with full bootstrap values in Fasta and Newick format in Supplementary Data.

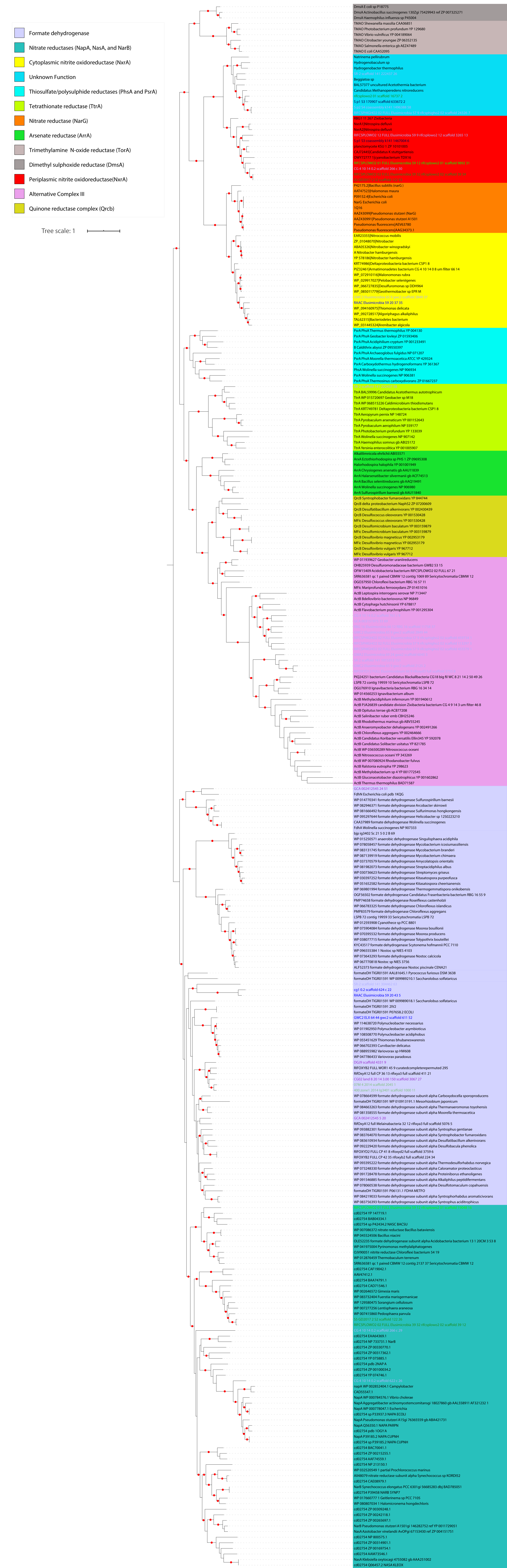
Formate dehydrogenase
Nitrate reductases (NapA, NasA, and NarB)
Cytoplasmic nitrite oxidoreductase (NxrA)
Unknown Function
Thiosulfate/polysulphide reductases (PhsA and PsrA)
Tetrathionate reductase (TtrA)
Nitrate reductase (NarG)
Arsenate reductase (ArrA)
Trimethylamine N-oxide reductase (TorA)
Dimethyl sulphoxide reductase (DmsA)
Periplasmic nitrite oxidoreductase(NxrA)
Alternative Complex III
Quinone reductase complex (Qrcb)
Tree scale: 1

Figure S12. Phylogenetic analysis of the dimethyl sulfoxide (DMSO) reductase superfamily (Maximum likelihood tree under a LG+F+G4 model of evolution). *Elusimicrobia* have an alternative complex III, a tetrathionate reductase (TtrA), cytoplasmic and periplasmic nitrite oxidoreductase (NXR), formate dehydrogenase and nitrate reductase (NapA and NasA). *Elusimicrobia* sequences are indicated with colored font. Scale bar indicates substitutions per site. The bootstraps are indicated by red circles when ≥ 85. The tree was made using sequences from (Castelle et al. 2013; Matheus Carnevali et al. 2019). The sequences and tree are available with full bootstrap values in Fasta and Newick format in Supplementary Data.

SUPPLEMENTARY DATASET

**Supplementary Dataset 1.** List of the 94 genomes used in this study. For each genome (column A), the NCBI genome accession, completeness and the contamination based on single copy genes are displayed in columns B, C and D respectively. Columns from E to J informs about the concatenated ribosomal proteins. Environments are provided in columns K and L, pubmed PMID in column M. The GTDB classification is shown in column N, classification based on 16S rRNA in column O. Final classification is shown in column P.

**Supplementary Dataset 2.** List of the 17 genomes from Myxococcales and Gracilibacteria genomes with the group VI nifH. Genome names and taxonomy are provided in columns A and C. When available, NCBI assembly accession is provided in column B.

**Supplementary Dataset 3.** Metabolic capacities across the 94 Elusimicrobia genomes. Genome names and environments are provided in columns A and B. Each metabolic feature is considered either complete (indicated by 1 and a red cell), partial (indicated by P and an orange cell) or absent (indicated by 0 and a white cell). Completeness of the pathways is based on literature and annotations (see the row 2 for partial information and the materials and methods for full details).
